# Sequential pathways preserve kinetochore integrity in meiosis

**DOI:** 10.64898/2026.11.05.764291

**Authors:** Marina Hamaia, Christos Spanos, Weronika Borek, Adele L. Marston

## Abstract

Kinetochores connect chromosomes to spindle microtubules, but how their integrity is maintained as they are remodelled during meiosis remains poorly understood. Here, we identify three distinct pathways linking centromeric chromatin to the outer kinetochore in budding yeast, mediated by the Ctf19 complex (Ctf19c^CCAN^), Mif2^CENP-C^ and binding of Okp1^CENP-Q^–Ame1^CENP-U^ to the Cse4^CENP-A^ essential N-terminal domain (END). Although partially redundant during mitosis, all three connections are required during meiosis, with distinct temporal requirements. The assembled Ctf19c^CCAN^ and Mif2^CENP-C^-dependent pathways maintain kinetochore integrity from meiotic entry, while Cse4^CENP-A^–Okp1^CENP-Q^–Ame1^CENP-U^ binding becomes critical during prophase and the meiotic divisions. We further show that Ipl1^Aurora^ ^B^ maintains kinetochore integrity independently of Ctf19c^CCAN^ and provide evidence for an outer kinetochore-associated Ipl1^Aurora^ ^B^ pool. Strikingly, Cse4^CENP-A^ phosphomimetic mutations that restore kinetochore integrity upon Ipl1^AuroraB^ depletion instead exacerbate defects in cells lacking Ctf19c^CCAN^. Our findings suggest that phosphorylation of the Cse4^CENP-A^ N-terminal domain (NTD) by Ipl1^Aurora^ ^B^ may regulate the use of alternative centromere-kinetochore connections to adapt to the changing requirements of meiosis.

## Introduction

The kinetochore is a large protein assembly that connects chromosomes to the spindle by linking centromeric CENP-A-containing chromatin to microtubules (Hinshaw & Harrison, 2018). In budding yeast, centromeres are marked by a single Cse4^CENP-A^- containing nucleosome (Camahort *et al*, 2007; Furuyama & Biggins, 2007; Greco *et al*, 2026). The histone chaperone Scm3^HJURP^ facilitates Cse4^CENP-A^ incorporation into chromatin through its interaction with the C-terminal centromere-targeting domain (CATD), before relocating to the essential N-terminal domain (END) to stabilise the nucleosome (Camahort *et al*, 2007; Shukla *et al*, 2024; Popchock *et al*, 2025). The Cse4^CENP-A^ nucleosome is recognised by the Mif2^CENP-C^ subunit and several components of the 13-subunit Ctf19 complex (Ctf19c, known as CCAN in humans), including Chl4^CENP-N^, Okp1^CENP-Q^ and Ame1^CENP-U^ (Fischböck-Halwachs *et al*, 2019; Yan *et al*, 2019; Dendooven *et al*, 2023). Interestingly, structural and biochemical studies have revealed a distinct mode of centromere recognition in which the Okp1^CENP-Q^- Ame1^CENP-U^ heterodimer binds directly to the END domain of Cse4^CENP-A^, independently of the remainder of the Ctf19c^CCAN^ (Carroll *et al*, 2009; Hornung *et al*, 2014; Xiao *et al*, 2017; Anedchenko *et al*, 2019; Hinshaw & Harrison, 2019; Dendooven *et al*, 2023; Deng *et al*, 2023). This interaction is incompatible with the architecture of the intact Cse4^CENP-A^-bound-Ctf19c^CCAN^, suggesting that Okp1^CENP-Q^- Ame1^CENP-U^ can establish an alternative connection between centromeric chromatin and the outer kinetochore. However, the functional significance of this alternative arrangement is unclear.

Okp1^CENP-Q^-Ame1^CENP-U^ form the COMA subcomplex, together with Ctf19^CENP-P^ and Mcm21^CENP-O^, but are the only Ctf19c^CCAN^ components, other than Mif2^CENP-C^ essential for mitotic growth (Ortiz *et al*, 1999; Ghosh *et al*, 2001; De Wulf *et al*, 2003; Fernius & Marston, 2009; Borek *et al*, 2021). Both Mif2^CENP-C^ and Ame1^CENP-U^ bind the Mtw1 complex (Mtw1c^MIS12c^, also known as MIND), which recruits the outer kinetochore Spc105^KNL1^ and Ndc80 complexes (Spc105c^KNL1c^ and Ndc80c^NDC80c^, respectively). Together, these complexes assemble into the KMN network which forms the microtubule-binding interface which is further stabilised by the Dam1 complex (Dam1c) (Hornung *et al*, 2014; Dimitrova *et al*, 2016; Hamilton *et al*, 2020). Cnn1^CENP-T^ provides an additional connection between Ctf19c^CCAN^ and Ndc80c^NDC80c^, although this pathway is dispensable for viability in *S. cerevisiae* (Malvezzi *et al*, 2013; Borek *et al*, 2021; Schleiffer *et al*, 2012). Thus, Okp1^CENP-Q^- Ame1^CENP-U^, along with Mif2^CENP-C^, provide the essential connections between centromeric chromatin and the outer kinetochore, while other Ctf19c^CCAN^ components, although dispensable for viability, contribute to chromosome segregation fidelity (De Wulf *et al*, 2003; Fernius & Marston, 2009).

Kinetochore assembly also requires Ipl1^Aurora^ ^B^, which phosphorylates Dsn1^DSN1^ to relieve autoinhibition of Mtw1c^MIS12c^ assembly (Akiyoshi *et al*, 2013; Dimitrova *et al*, 2016; Bonner *et al*, 2019; Killinger *et al*, 2020; Turner *et al*, 2026) and Cse4^CENP-A^ to promote its interaction with Scm3^HJURP^ (Popchock *et al*, 2025). Several pathways have been implicated in targeting the Ipl1-containing chromosome passenger complex (CPC) to centromeres in mitosis through interactions of its Bir1^SURVIVIN^ subunit with Ndc10 (Yoon & Carbon, 1999), Sgo1 or histone H3S3ph (Dai *et al*, 2005; Wang *et al*, 2010). Surprisingly, deletion of the N-terminal 228 amino acids of CPC component Sli15 (*sli15ΔN*), which removes the CEN box required for Bir1^SURVIVIN^ interaction and thus centromere targeting, is tolerated in both mitosis and meiosis, with near-normal chromosome segregation fidelity. Moreover, Ipl1^Aurora^ ^B^ localisation appears unaffected in mitosis (Campbell & Desai, 2013). Bir1-independent CPC recruitment to inner kinetochores via Ctf19^CENP-P^ (Fischböck-Halwachs *et al*, 2019; García-Rodríguez *et al*, 2019) or to outer kinetochores via Spc105-Kre28^KNL1c^ (Dudziak *et al*, 2025) have been proposed to explain these observations.

The specialised meiotic cell cycle imposes additional demands on kinetochores. Following DNA replication, during the extended meiotic prophase, the outer kinetochore disassembles before being reassembled for two distinct and successive chromosome segregation events. During meiosis I, sister kinetochores co-orient towards the same pole to allow homologous chromosomes to segregate, while sister chromatids segregate during meiosis II. Intriguingly, although most Ctf19c^CCAN^ subunits are dispensable for mitotic growth, they become critical during meiosis, with their loss severely compromising kinetochore integrity, spore formation and spore viability (Ghosh *et al*, 2001; Borek *et al*, 2021). Notable exceptions are Nkp1 and Nkp2, which have relatively minor roles in meiosis (Fernius & Marston, 2009). Why meiotic kinetochores are particularly dependent on Ctf19c^CCAN^, and whether this reflects the proposed role of Ctf19^CENP-P^ in recruiting Ipl1^Aurora^ ^B^ remains unknown.

Here, we identify three pathways linking centromeric chromatin to the outer kinetochore that are individually required during meiosis but are redundant in mitosis: (i) Mif2^CENP-C^, (ii) the Ctf19c^CCAN^; and (iii) direct binding of Okp1^CENP-Q^- Ame1^CENP-U^ to the Cse4^CENP-A^ END. We demonstrate that Mif2^CENP-C^ and Ctf19c^CCAN^ are specifically required to maintain kinetochore integrity until exit from prophase I, while direct binding of Okp1^CENP-Q^- Ame1^CENP-U^ to the Cse4^CENP-A^ END is required during the meiotic divisions. Ipl1^Aurora^ ^B^ is required throughout meiosis and acts independently of Ctf19c^CCAN^, localising instead at the outer kinetochore. Our findings suggest a model in which dynamic phosphorylation of Cse4^CENP-A^ NTD by Ipl1^Aurora^ ^B^ coordinates the sequential activation of recruitment pathways to maintain kinetochore integrity throughout meiosis.

## Results

### Ctf19^CENP-P^ maintains kinetochore integrity from meiotic entry until prophase I exit

To reveal the precise timing and duration of the role of Ctf19^CENP-P^ in kinetochore assembly in meiosis, we used an auxin-inducible degron to acutely deplete Ctf19^CENP-P^ at different meiotic stages and assessed Mtw1^MIS12^-tdTomato signal intensity as a readout of kinetochore integrity. Cells were arrested at meiotic entry (G1) and synchronously released into meiosis (Berchowitz *et al*, 2013). *CTF19^CENP-P^-3V5-AID* cells were mock-treated with DMSO or treated with 1-Naphthaleneacetic acid (NAA), a synthetic auxin analogue, from 2 hours before the release, at the release, and 2 or 4 hours after the release (Figure 1A). Western blotting confirmed robust depletion of Ctf19^CENP-P^-3V5-AID within 30 minutes of auxin addition, while the presence of the meiosis-specific cohesin Rec8 and completion of DNA replication confirmed entry into meiosis (Figure 1B; Figure S1A-E). Degradation of Ctf19^CENP-P^-3V5-AID either 2 hours before, at, or 2 hours after meiotic entry caused a substantial and sustained decrease in the percentage of cells with bright Mtw1^MIS12^-tdTomato signal (Figure 1C). In contrast, depleting Ctf19^CENP-P^-3V5-AID from 4 hours after meiotic entry had a much milder effect, with a similar proportion of cells having a bright Mtw1^MIS12^-tdTomato signal as the mock-treated DMSO control (Figure 1C). Therefore, towards the end of prophase I, cells become desensitised to the loss of Ctf19^CENP-P^.

**Figure 1.**
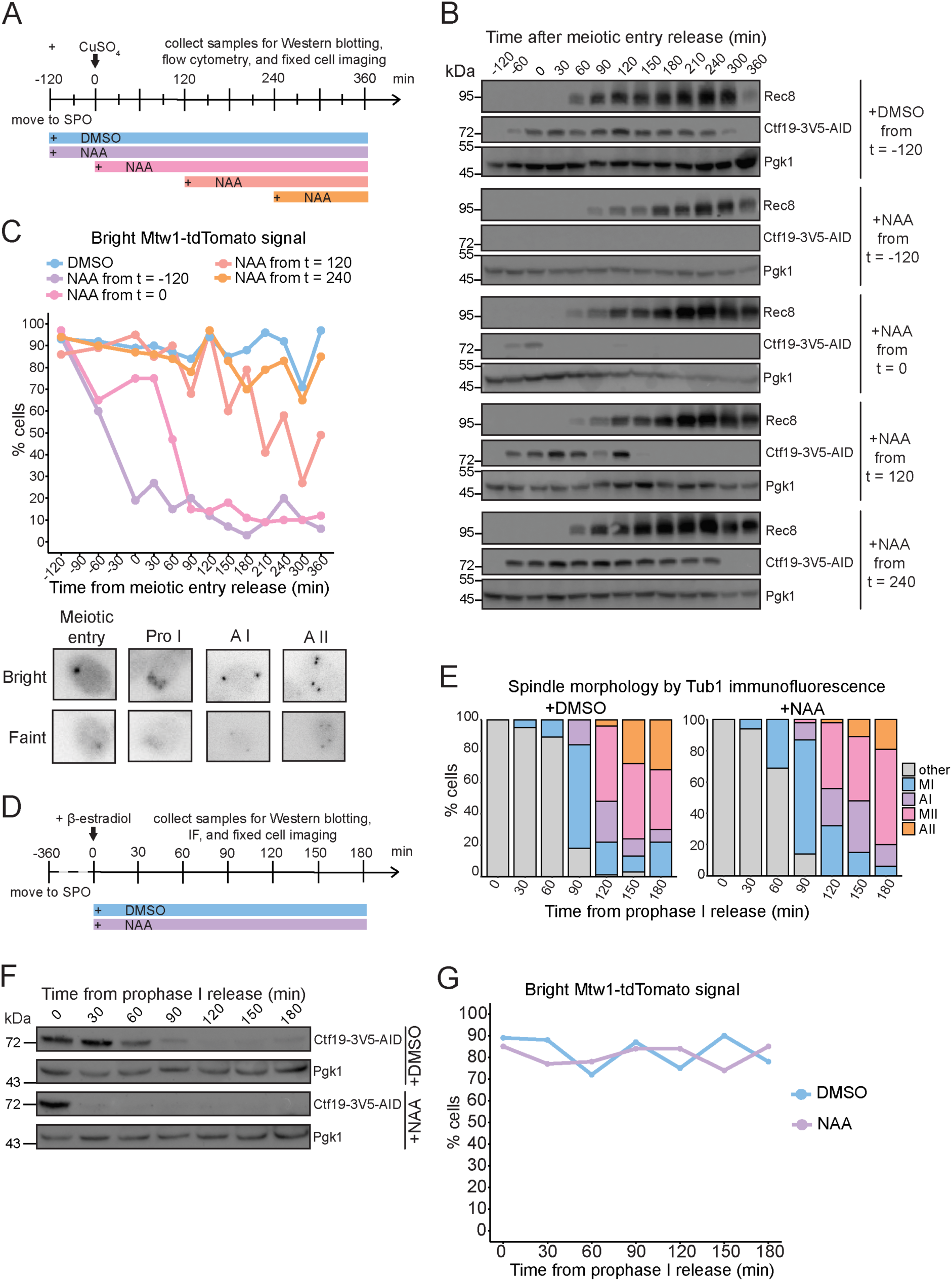
Ctf19^CENP-P^ maintains kinetochore integrity from meiotic entry until prophase I exit. (A) Schematic of the experimental timeline. Diploid cells expressing *ctf19-3V5-AID* and the *pCUP-IME1 pCUP-IME4* system were induced to undergo meiosis. After 2 h, copper (II) sulfate was added to release cells synchronously into meiotic entry. DMSO or 1-Naphthylacetic acid (NAA) was added to the cells from 2 h before the release, from the release, from 2 h after the release, or from 4 h after release. Samples for Western blotting, flow cytometry, and fixed cell imaging were taken every 30 min. (B) Degradation of Ctf19-3V5-AID upon NAA addition. Western blotting of samples collected in (A) with α-Rec8, α-V5 (Ctf19), and α-Pgk1. (C) Depletion of Ctf19-3V5-AID in early meiosis causes loss of kinetochore integrity. Percentage of cells with ‘bright’ Mtw1-tdTomato signal in fixed cell samples from timecourse in (A). Inset: representative images of ‘bright’ and ‘faint’ Mtw1-tdTomato signal at meiotic entry, prophase I, anaphase I, and anaphase II of meiosis. n = 100 cells scored at each timepoint. (D) Schematic of the experimental timeline. Diploid cells expressing *ctf19-3V5-AID* and the Ndt80 prophase I arrest-release system were induced to undergo meiosis. After 6 h, β-estradiol was added to release cells synchronously from prophase I. DMSO or NAA was added at release. Samples for Western blotting, immunofluorescence, and fixed cell imaging were taken every 30 min. (E) Meiosis progression is unaffected by Ctf19-3V5-AID depletion after prophase I exit. Quantification of meiotic stage of 100 cells at each timepoint of samples collected in (D) based on Tub1 immunofluorescence. n = 200 cells scored at each timepoint. (F) Degradation of Ctf19-3V5-AID upon NAA addition. Western blotting of samples collected in (D) with α-V5 (Ctf19) and α-Pgk1. (G) Depletion of Ctf19-3V5-AID after prophase I exit does not affect kinetochore integrity. Percentage of cells with ‘bright Mtw1-tdTomato signal in fixed cell samples from timecourse in (D). n = 100 cells scored at each timepoint.

To test whether cells require Ctf19^CENP-P^ for kinetochore integrity beyond prophase I, cells were synchronously released from prophase arrest (Carlile & Amon, 2008). Ctf19^CENP-P^-3V5-AID cells were either mock-treated with DMSO or degraded with auxin (Figure 1D). Progression through the meiotic divisions was determined by spindle morphology, and no substantial delay was detected when Ctf19^CENP-P^-3V5-AID was depleted, which was confirmed by immunoblotting (Figure 1E-F). Interestingly, Ctf19^CENP-P^-3V5-AID levels greatly decreased in the DMSO control 2 hours after prophase I release (Figure 1E-F). Similarly, Ctf19^CENP-P^-3V5-AID levels decreased at later timepoints in the *pCUP-IME1 pCUP-IME4* system (Figure 1B). The spore viability of *CTF19 ^CENP-P^-3V5-AID* cells was similar to the wild type (Figure S1F), suggesting that kinetochore assembly during the meiotic divisions requires only a small amount of Ctf19^CENP-P^. Nevertheless, depletion of Ctf19^CENP-P^-3V5-AID from prophase I exit had no effect on Mtw1^MIS12^-tdTomato signal intensity at kinetochores (Figure 1G). We conclude that Ctf19^CENP-P^ is essential for maintaining kinetochore integrity in meiosis from meiotic entry until prophase I exit, whereupon kinetochores are no longer dependent on Ctf19^CENP-P^.

### Ipl1^Aurora^ ^B^ is needed to maintain kinetochore integrity throughout meiosis

We sought to determine whether the essential role of Ctf19^CENP-P^ in meiotic kinetochore assembly is to recruit the CPC. To deplete Ipl1^Aurora^ ^B^ in meiosis, we used the *pSCC1-IPL1* allele, which places *IPL1^Aurora^ ^B^* expression under the control of the mitosis-specific *SCC1* promoter (Monje-Casas *et al*, 2007). In addition to the kinetochore label, Mtw1^MIS12^-tdTomato, cells also carried Zip1-GFP and Tub1-GFP to label the synaptonemal complex and spindle, respectively, and thereby assess the meiotic stage. Mtw1^MIS12^-tdTomato live imaging showed defective dispersal of centromeres from the RabI cluster in *pSCC1-IPL1^Aurora^ ^B^*, as expected (Meyer *et al*, 2013). Importantly, a significant decrease in Mtw1^MIS12^-tdTomato signal intensity was observed from prophase I exit onwards, as determined by loss of Zip1-GFP signal (Figure 2A-B). Strikingly, this is the stage of meiosis when Ctf19^CENP-P^ is no longer required for kinetochore integrity. However, it remained possible that the *pSCC1-IPL1^Aurora^ ^B^*allele allowed residual Ipl1^Aurora^ ^B^ to persist in early meiosis, precluding a firm conclusion about the requirement of Ipl1^Aurora^ ^B^ in kinetochore integrity at meiotic entry. To overcome this, we tagged the CPC component Sli15 with an auxin-inducible degron and directly compared Mtw1^MIS12^-tdTomato signal in *SLI15-3V5-AID* or *CTF19^CENP-P^-3V5-AID* diploids upon treatment with auxin or DMSO from 2 hours before synchronous release into meiotic entry (Berchowitz *et al*, 2013) (Figure 2C). Robust depletion of Ctf19*^CENP-P^*-3V5-AID and Sli15-3V5-AID and completion of DNA replication were monitored by immunoblotting (Figure 2D) and flow cytometry (data not shown), respectively. Quantification of the Mtw1^MIS12^-tdTomato signal intensity showed that degrading Sli15-3V5-AID caused a reduction in Mtw1^MIS12^-tdTomato signal from before meiotic entry and throughout meiosis, compared to the DMSO-treated control (Figure 2E). This was similar to the decrease in Mtw1^MIS12^-tdTomato signal observed in Ctf19*^CENP-P^*-3V5-AID cells treated with auxin (Figure 2F) and contradictory to the live imaging of *pSCC1-IPL1^Aurora^ ^B^*cells (Figure 2B). This discrepancy may be attributed to residual levels of Ipl1^Aurora^ ^B^ activity being present in early meiosis in the *pSCC1-IPL1^Aurora^ ^B^* mutant, which may be sufficient to support kinetochore assembly. The Mtw1^MIS12^-tdTomato kinetochore defect was specific to loss of Ctf19^CENP-P^ or Ipl1^Aurora^ ^B^ because inhibiting the kinetochore-associated checkpoint kinase Mps1 in meiosis did not affect kinetochore integrity (Figure S2A-B). Therefore, Sli15^INCENP^-Ipl1^Aurora^ ^B^ is required for kinetochore integrity from meiotic entry, similar to Ctf19^CENP-P^.

**Figure 2.**
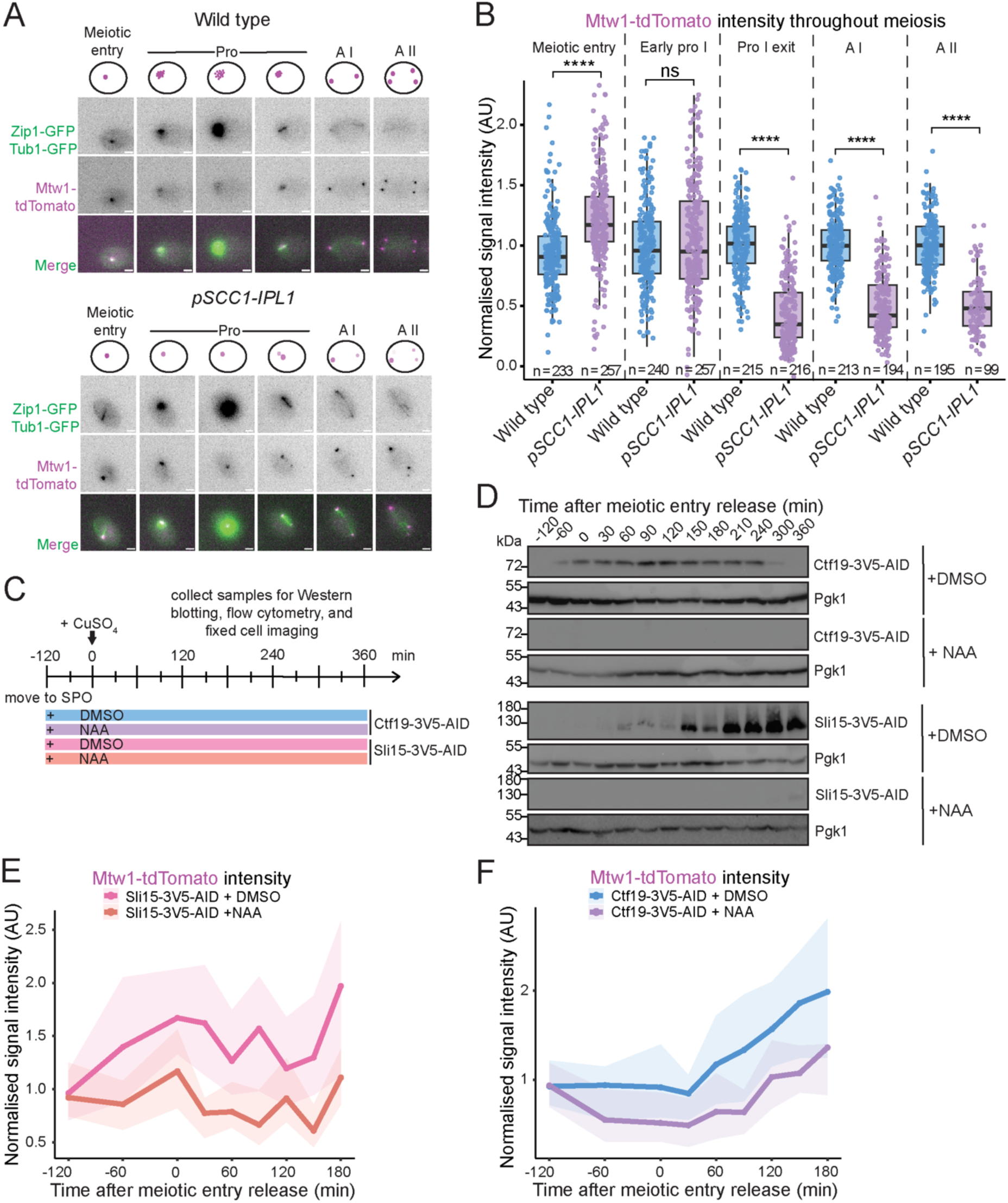
Ipl1^Aurora^ ^B^ is needed to maintain kinetochore integrity throughout meiosis. (A) Representative images from live imaging of wild type and *pSCC1-IPL1* diploids with Zip1-GFP, Tub1-GFP, and Mtw1-tdTomato. All scale bars represent 2 μm. (B) *pSCC1-IPL1* cells have a kinetochore defect from prophase I exit. Quantification of normalised Mtw1-tdTomato signal intensity throughout meiosis. Meiotic stage was determined based on Zip1-GFP and Tub1-GFP signal and morphology. Data shown are from 3 biological replicates. Boxplots represent interquartile ranges (IQR), and the middle line is the median. Whiskers extend to the furthest data points within 1.5 IQR from the 25th and 75th percentiles; data points outside this range are plotted individually as outliers; Mann-Whitney test. (C) Schematic of experimental timeline. Diploid cells expressing either *ctf19-3V5-AID* or *sli15-3V5-AID* and the *pCUP-IME1 pCUP-IME4* system were induced to undergo meiosis. DMSO or NAA was added to the cells. After 2 h, copper (II) sulfate was added to release cells synchronously into meiotic entry. Samples for Western blotting, flow cytometry, and fixed cell imaging were taken every 30 min. (D) Degradation of Ctf19-3V5-AID and Sli15-3V5-AID upon NAA addition. Western blotting of samples collected in (C) with α-V5 (Ctf19 or Sli15), and α-Pgk1. (E) Depletion of Sli15-3V5-AID in early meiosis causes loss of kinetochore integrity. Normalised signal intensity of Mtw1-tdTomato in *sli15-3V5-AID* cells treated with DMSO or NAA. n = 100 cells were quantified at each timepoint. (F) Depletion of Ctf19-3V5-AID in early meiosis causes loss of kinetochore integrity. Normalised signal intensity of Mtw1-tdTomato in *ctf19-3V5-AID* cells treated with DMSO or NAA. n = 100 cells were quantified at each timepoint.

### The outer kinetochore, rather than Ctf19, mediates Bir1-independent CPC recruitment during meiosis

Ctf19^CENP-P^ has been implicated in recruitment of the CPC to kinetochores in mitosis (Fischböck-Halwachs *et al*, 2019; García-Rodríguez *et al*, 2019). One explanation for our findings is that Ctf19^CENP-P^ recruits Ipl1^Aurora^ ^B^ early in meiosis, with Ctf19^CENP-P^-independent pathways maintaining Ipl1^Aurora^ ^B^ recruitment after prophase exit. To test this idea, we sought to understand which pathways recruit Ipl1^Aurora^ ^B^ to kinetochores at different stages in meiosis. Ipl1^Aurora^ ^B^ can be recruited either through Bir1-dependent pathways, which depend on Sli15’s N-terminal region, or Bir1-independent pathways, which include recruitment to outer kinetochores through interaction with Spc105-Kre28, or the proposed inner kinetochore recruitment through Ctf19^CENP-P^ interaction (Figure 3A). Abolition of the Bir1-dependent pathways using the *sli15ΔN* mutation caused no significant difference in Mtw1^MIS12^-tdTomato intensity or associated Ipl1^Aurora^ ^B^-mNeonGreen signal compared to wild type (Figure 3B-D; Figure S3A-B). Consistently, *sli15ΔN* has no spore viability defect (Figure S3C; (Campbell & Desai, 2013)) and deletion of the Haspin kinase orthologues, *ALK1* and *ALK2*, also has no effect on Ipl1^Aurora^ ^B^-mNeonGreen co-localisation with Mtw1^MIS12^-tdTomato (Figure S3D-F). Therefore, Bir1-dependent centromeric targeting of Ipl1^Aurora^ ^B^ is not necessary for meiotic kinetochore assembly.

**Figure 3.**
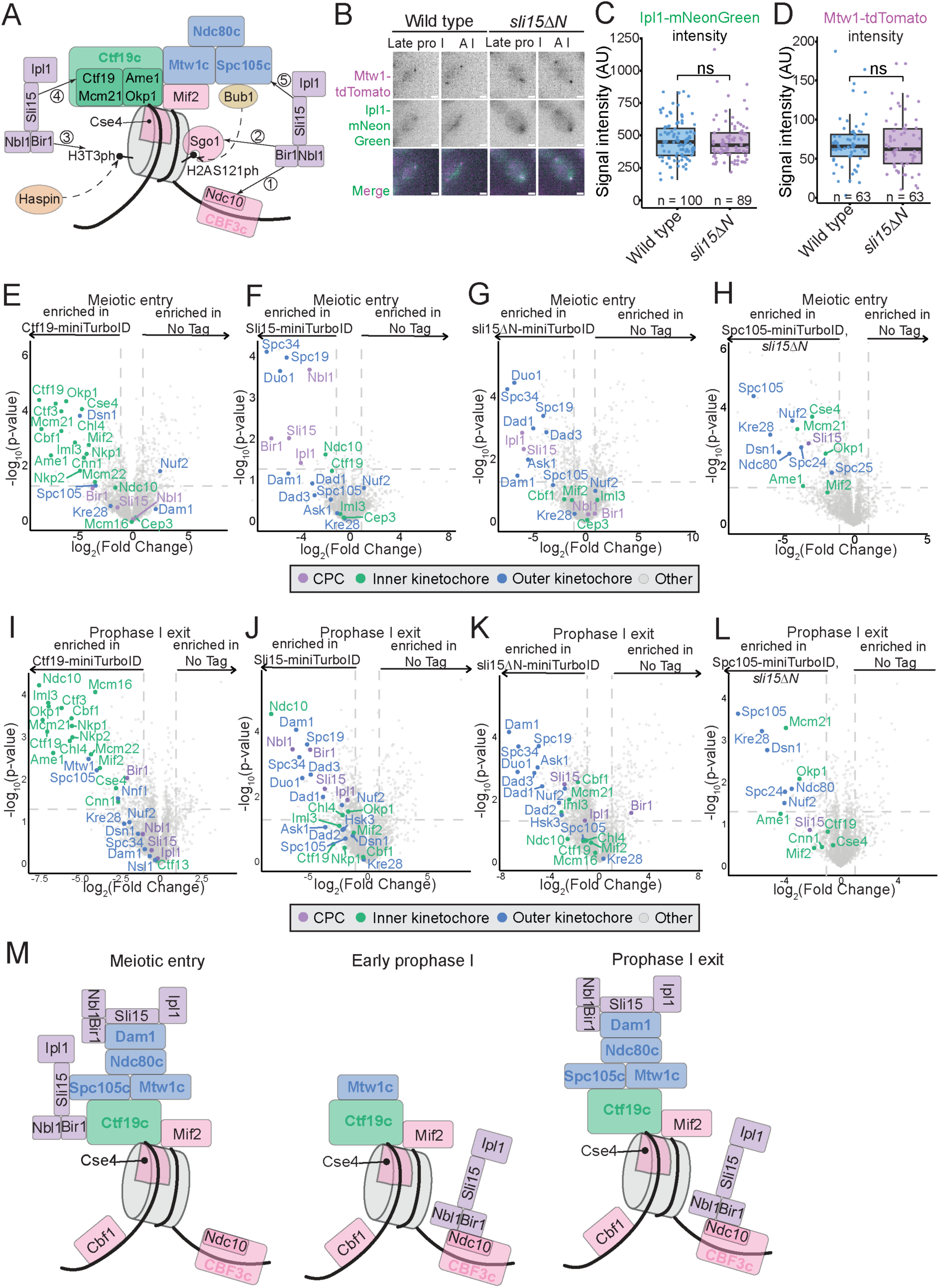
The outer kinetochore, rather than Ctf19, mediates Bir1-independent CPC recruitment during meiosis. (A) Schematic of CPC recruitment pathways to centromeres and kinetochores in mitosis; (1) Bir1 interacts with Ndc10; (2) Bub1 phosphorylation of H2AS121 recruits Sgo1 which interacts with Bir1; (3) Haspin kinase phosphorylates H3T3 which recruits Bir1; (4) Sli15 interaction with Ctf19; (5) Sli15 interaction with Spc105 and Kre28. (B) Representative images from live imaging of Mtw1-tdTomato and Ipl1-mNeonGreen at late prophase I and anaphase I in wild type and *sli15ΔN* cells. All scale bars represent 2 μm. (C-D) Absence of Bir1-dependent CPC recruitment shows no Ipl1 localisation or outer kinetochore assembly defect in meiosis. (C) Quantification of Ipl1-mNeonGreen signal intensity in (B) at late prophase I. (D) Absence of Bir1-dependent CPC recruitment shows no outer kinetochore assembly defect in meiosis. Quantification of Mtw1-tdTomato signal intensity in (B) at late prophase I. Boxplots represent interquartile ranges (IQR), and the middle line is the median. Whiskers extend to the furthest data points within 1.5 IQR from the 25th and 75th percentiles; data points outside this range are plotted individually as outliers; Mann-Whitney test. (E-H) The CPC is in close proximity to the outer kinetochore rather than Ctf19 at meiotic entry. Volcano plots presenting the mass spectrometry-identified biotinylated proteins pulled down by streptavidin-coated beads at meiotic entry compared to no tag in diploids expressing (E) Ctf19-miniTurboID; (F) Sli15-miniTurboID; (G) sli15ΔN-miniTurboID; (H) Spc105-miniTurboID *sli15ΔN*. Diploids containing the *pCUP-IME1 pCUP-IME4* system were induced to undergo meiosis and harvested after 2 h without release from meiotic entry arrest. Data are from 3 biological replicates. Log2(fold change) between conditions is shown with corresponding p-values. Dashed line indicates log_2_(fold change) = |2|. (I-L) The CPC is in close proximity to the Dam1 complex rather than Ctf19 at prophase I exit. Volcano plots presenting the mass spectrometry-identified biotinylated proteins pulled down by streptavidin-coated beads at prophase I exit compared to no tag in diploids expressing (E) Ctf19-miniTurboID; (F) Sli15-miniTurboID; (G) sli15ΔN-miniTurboID; (H) Spc105-miniTurboID *sli15ΔN*. Diploids containing the Ndt80 block-release system were induced to undergo meiosis and released from arrest by β-estradiol addition at 6 h. Cells were harvested 30 min later. Data are from 3 biological replicates. Log2(fold change) between conditions is shown with corresponding p-values. Dashed line indicates log_2_(fold change) = |2|. (M) Model of CPC localisation throughout meiosis.

To understand which other pathways might recruit Ipl1^Aurora^ ^B^ to kinetochores, we took a proximity-dependent in vivo biotinylation approach (Branon *et al*, 2018; Sears *et al*, 2019). Cells expressing either Ctf19^CENP-P^-miniTurboID-3V5, Sli15^INCENP^-miniTurboID-3V5, sli15^INCENP^ΔN-miniTurboID-3V5, or Spc105^KNL1^-miniTurboID-3V5 were arrested at meiotic entry, a time when both Ctf19^CENP-P^ and Ipl1^Aurora^ ^B^ are important for kinetochore assembly and were *in vivo* labelled with biotin for 50 minutes prior to harvesting. Biotinylated proteins were isolated using streptavidin-coated beads and identified by mass spectrometry. As expected, inner kinetochore subunits, including Ctf19^CENP-P^, were highly enriched in the Ctf19^CENP-P^-miniTurboID-3V5 samples compared to a no tag control (Figure 3E); however, CPC subunits were not. Similarly, Sli15^INCENP^-miniTurboID-3V5 samples were enriched for CPC subunits but showed little enrichment of the inner kinetochore, with Ctf19^CENP-P^ itself not significantly enriched. The only enriched inner kinetochore protein was Ndc10 of the Cbf3 complex; however, this pathway of Ipl1^Aurora^ ^B^ recruitment is Bir1-dependent and, accordingly, Ndc10 was absent from the sli15^INCENP^ΔN-miniTurboID-3V5 samples. Instead, strikingly, the outer kinetochore was highly enriched in both the Sli15^INCENP^-miniTurboID-3V5 and sli15^INCENP^ΔN-miniTurboID-3V5 samples, particularly subunits of the Dam1 complex (Duo1, Spc34, and Spc19; Figure 3F-G). The Spc105^KNL1^-miniTurboID-3V5 *sli15^INCENP^ΔN* samples also showed strong enrichment of both inner and outer kinetochore proteins over no tag, and, notably, Sli15^INCENP^ was also significantly enriched (Figure 3H). Therefore, we find no evidence that Ctf19^CENP-P^ recruits CPC during meiosis. Instead, our findings support a model in which Spc105^KNL1^ and the Dam1c mediate Bir1-independent CPC recruitment at meiotic entry.

We additionally examined pathways of Ipl1^Aurora^ ^B^ localisation at prophase I exit, when kinetochores become desensitised to Ctf19^CENP-P^ loss (Figure 1D-G). At prophase I exit, inner kinetochore subunits were highly enriched in Ctf19^CENP-P^-miniTurboID-3V5 samples over no tag (Figure 3I; S3G-H). Bir1^SURVIVIN^ was also significantly enriched, but this enrichment was lost in Ctf19^CENP-P^-miniTurboID-3V5 *sli15^INCENP^ΔN* samples, suggesting that Ctf19^CENP-P^ is proximal to the Bir1^SURVIVIN^-dependent centromere pool, rather than the Bir1^SURVIVIN^-independent pathway required for kinetochore assembly (Figure S3G-H). Furthermore, Ctf19^CENP-P^ was not detected at significant levels in either Sli15^INCENP^-miniTurboID-3V5 or sli15^INCENP^ΔN-miniTurboID-3V5 samples (Figure 3J-K). Notably, Ndc10 was highly enriched in Sli15^INCENP^-miniTurboID-3V5, but not in Sli15^INCENP^ΔN-miniTurboID-3V5 samples, suggesting increased centromere recruitment at prophase exit compared to at meiotic entry (Figure 3J-K; S3I). The outer kinetochore Dam1 complex subunits were again highly enriched in these samples (Figure 3J-K; S3I). However, in contrast to meiotic entry, Sli15^INCENP^ was not significantly enriched in Spc105^KNL1^-miniTurboID-3V5 *sli15^INCENP^ΔN* samples at prophase exit (Figure 3L). This could be explained by shedding of the outer kinetochore in early prophase, leading to loss of this pool and re-establishment of only the Dam1-associated Ipl1^Aurora^ ^B^ pool following outer kinetochore reassembly at prophase exit (Figure 3M). In summary, we find no evidence that Ctf19^CENP-P^ recruits Ipl1^Aurora^ ^B^ to the inner kinetochore at either meiotic entry or prophase exit. Taken together with the distinct temporal requirements for Ipl1^Aurora^ ^B^ and Ctf19^CENP-P^ in maintenance of kinetochore integrity, these findings support the conclusion that they act in separate pathways.

### Ipl1^Aurora^ ^B^ promotes meiotic kinetochore assembly by phosphorylating Cse4^CENP-A^ and Dsn1

Next, we addressed how Ipl1^Aurora^ ^B^ might promote kinetochore assembly in meiosis. In mitotic cells, Ipl1^Aurora^ ^B^ phosphorylates the Cse4^CENP-A^ END at Ser40, which promotes Scm3^HJURP^ binding to the Cse4^CENP-A^ END and the localisation of Cse4^CENP-A^ at the centromeric nucleosome ((Figure 4A); (Popchock *et al*, 2025)). Additionally, Ipl1^Aurora^ ^B^ phosphorylates the outer kinetochore protein Dsn1 at Ser240 and Ser250, relieving Mtw1^MIS12^ complex autoinhibition and promoting outer kinetochore association with the inner kinetochore during mitosis (Akiyoshi *et al*, 2013; Dimitrova *et al*, 2016; Turner *et al*, 2026). We asked whether phosphomimetic mutants of Cse4^CENP-A^ and Dsn1 could rescue defective meiotic kinetochore assembly in the absence of Ipl1^Aurora^ ^B^. We confirmed that, in an otherwise wild-type background, spore viability was not greatly affected by various combinations of phosphomimetic mutants in the Cse4^CENP-A^ NTD domain, though *cse4^CENP-A^-40D* alone resulted in a modest increase in tetrads with one inviable spore (Figure 4B). Nevertheless, consistent with mitotic studies (Popchock *et al*, 2025), *cse4^CENP-A^-S40D* significantly increased Mtw1^MIS12^-tdTomato signal intensity from meiotic entry in *pSCC1-IPL1^Aurora^ ^B^* cells. Intriguingly, *cse4^CENP-A^-S22D* also significantly rescued Mtw1^MIS12^-tdTomato signal intensity, but only from early prophase. By prophase exit, where defective kinetochore assembly is most apparent in *pSCC1-IPL1^Aurora^* ^B^, each of these phosphomimetic mutants, including the double *cse4^CENP-A^-S22D,* S*40D* (*cse4-2D*) mutant, showed a substantial rescue of Mtw1^MIS12^-tdTomato signal (Figure 4B-F). The mean prophase I duration in *pSCC1-IPL1^Aurora^ ^B^* was significantly shorter than wild type (249 min vs. 304 min), whereas both *cse4^CENP-A^-S22D* and *cse4^CENP-A^-S40D* extended prophase in *pSCC1-IPL1^Aurora^ ^B^* cells to 353 min and 335 min, respectively (Figure 4G). Although the basis for this timing difference is unclear, the additional time may facilitate complete kinetochore assembly and thereby contribute to the rescue. Consistent with *cse4^CENP-A^-S40D* improving functional kinetochore assembly *in pSCC1-IPL1^Aurora^ ^B^* cells, we found that correct segregation of homozygously GFP-labelled *CENV* during meiosis I increased from 22% in *pSCC1-IPL1^Aurora^ ^B^* cells to 40% in *pSCC1-IPL1^Aurora B^ cse4^CENP-A^-S40D* cells (Figure 4H). Therefore, Ipl1^AuroraB^ promotes accurate chromosome segregation in meiosis, in part through directing proper kinetochore assembly.

**Figure 4.**
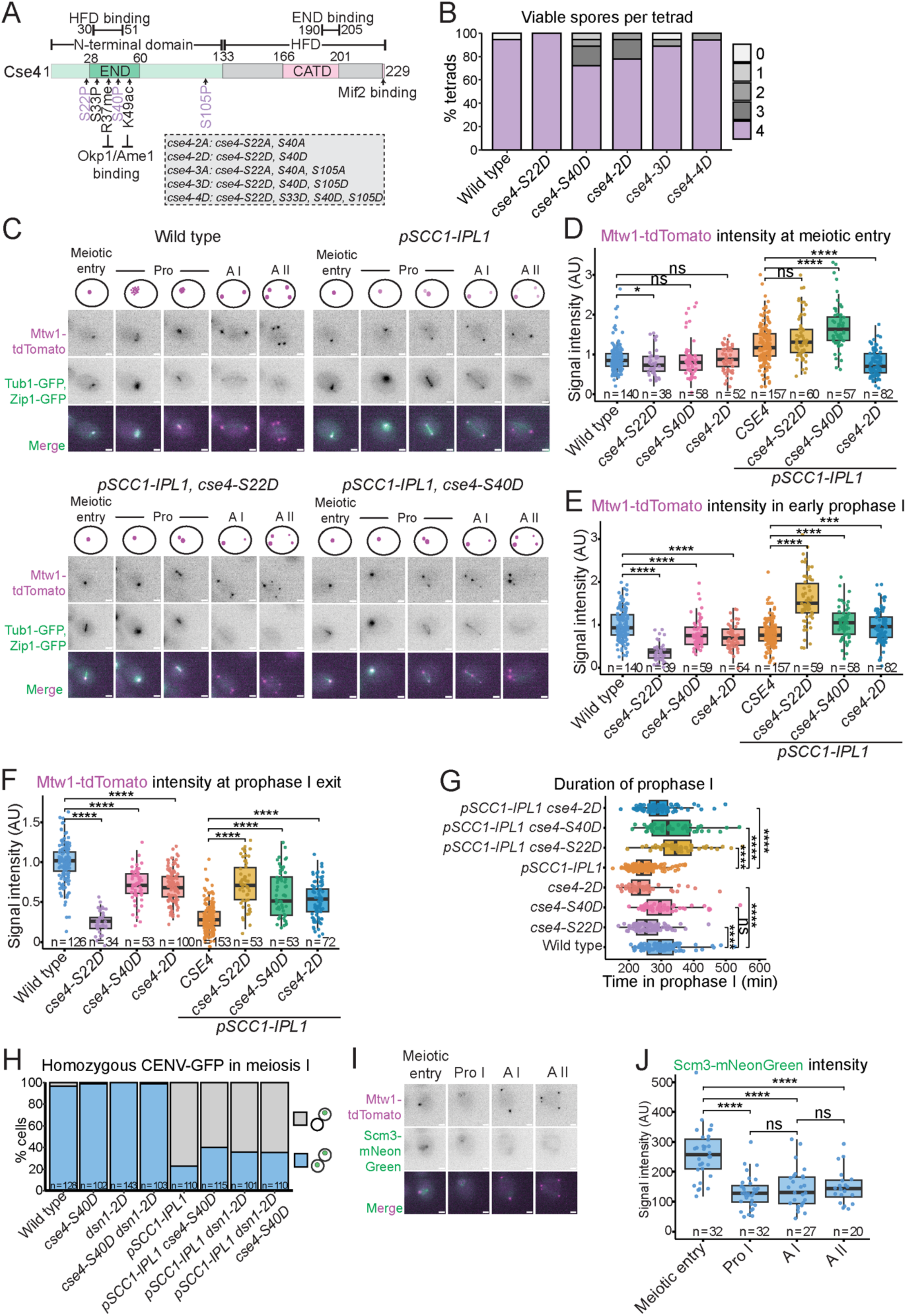
Ipl1^Aurora^ ^B^ promotes meiotic kinetochore assembly by phosphorylating Cse4^CENP-A^ and Dsn1. (A) Schematic of Cse4 protein with domains, regions interacting with Mif2^CENP-C^ and Okp1^CENP-Q^-Ame1^CENP-U^, and post-translationally modified residues. Ipl1^Aurora^ ^B^ target sites are highlighted in lilac. Inset: point mutations in the *cse4* mutant alleles used in this study. (B) Phosphomimetic mutations of *Cse4* do not affect spore viability. The number of viable progeny was scored 48 h after tetrad dissection. n = 18 tetrads. (C) Representative images, and illustrative schematics above, of live imaging of Mtw1-tdTomato, Tub1-GFP, and Zip1-GFP in meiosis. All scale bars represent 2 μm. (D-F) *cse4-S22D* and *cse4-S40D* rescue the *pSCC1-IPL1* kinetochore defect in meiosis. Quantification of Mtw1-tdTomato signal intensity at (D) meiotic entry; (E) early prophase I; (F) prophase I exit. Data are from 2 biological replicates. (G) Cse4 phosphomimetic mutations increase the length of prophase I in *pSCC1-IPL1* cells. Prophase I duration was measured between the appearance and disappearance of Zip1-GFP signal in live imaging shown in (C). Data are from 2 biological replicates. (H) *cse4-S40D* and *dsn1-2D (dsn1-S240D, S250D)* mildly improve meiosis I segregation defect of *pSCC1-IPL1* cells. The percentage of binucleate cells with both copies of chromosome V marked with GFP at the *URA3* locus with the indicated GFP dot segregation was determined. n > 100 binucleates for each strain. (I) Representative images from live imaging of Mtw1-tdTomato and Scm3-mNeonGreen in meiosis. All scale bars represent 2 μm. (J) Scm3 kinetochore localisation decreases in prophase I. Quantification of Scm3-mNeonGreen signal intensity from (I). Meiotic stage determined by Mtw1-tdTomato foci morphology. In (D-G) and (J), boxplots represent interquartile ranges (IQR), and the middle line is the median. Whiskers extend to the furthest data points within 1.5 IQR from the 25th and 75th percentiles; data points outside this range are plotted individually as outliers; Mann-Whitney test.

We next tested whether phosphomimetic mutations in the other known Ipl1^Aurora B^ target required for kinetochore assembly, Dsn1, further improved kinetochore function in *pSCC1-IPL1^Aurora^ ^B^* cells. Although *pSCC1-IPL1^Aurora^ ^B^ dsn1-S240, S250D* cells showed improved segregation fidelity (from 22% to 36%), combining the Cse4^CENP-A^ and Dsn1 phosphomimetic residues did not provide further rescue, with 35% of *pSCC1-IPL1^Aurora^ ^B^ dsn1-S240, S250D cse4^CENP-A^-S40D* cells correctly segregating homozygous *CENV-*GFP foci (Figure 4H). Because chromosome segregation reflects both Ipl1^Aurora^ ^B^-dependent kinetochore assembly and error correction, the lack of additional rescue may be limited by persistent defects in error correction, masking any further improvement in kinetochore assembly.

Ipl1^Aurora^ ^B^-dependent phosphorylation of Cse4^CENP-A^-S40 promotes binding of the Scm3^HJURP^ chaperone to promote Cse4^CENP-A^ deposition (Popchock *et al*, 2025). Our finding that Cse4^CENP-A^-S40D enhances Mtw1^MIS12^-tdTomato levels in *pSCC1-IPL1^Aurora^ ^B^* from meiotic entry (Figure 4C-F) suggested that Scm3^HJURP^ may localise to kinetochores at this time. Indeed, a strong Scm3^HJURP^-mNeonGreen signal was detected at meiotic entry; however, signal intensity declined significantly from prophase I and remained low through anaphase II (Figure 4I-J). These results suggest that phosphorylation-dependent association of Scm3^HJURP^ with the Cse4^CENP-A^ N-terminal domain stabilises kinetochores specifically during early meiosis. We conclude that Ipl1^Aurora^ ^B^ stabilises kinetochores through at least two mechanisms during meiosis: by promoting the interaction between the inner and outer kinetochore via Dsn1^DSN1^ phosphorylation, and by stabilising the inner kinetochore at centromeres through promoting Scm3^HJURP^ association with Cse4^CENP-A^. The latter mechanism is restricted to early meiosis.

### Cse4 phosphorylation precludes kinetochore assembly in *ctf19Δ* cells

Our findings suggest that Ctf19^CENP-P^ maintains kinetochore integrity through a distinct pathway to Ipl1^Aurora^ ^B^ (Figure 3D-F; 3H-J). Nevertheless, it remained possible that the two pathways could compensate for each other in meiosis. We therefore tested whether the Cse4^CENP-A^ phosphomimetic mutants that rescue the *pSCC1-IPL1^Aurora^ ^B^* kinetochore defect could also rescue the *ctf19Δ* kinetochore defect. Surprisingly, however, the Cse4^CENP-A^ phosphomimetic mutations not only failed to rescue the *ctf19Δ* phenotype but prevented these cells from sporulating (Figure 5A). Consistently, Mtw1^MIS12^-tdTomato signal intensity was not rescued in *ctf19Δ* cells by any of the Cse4^CENP-A^ phosphomimetic mutants and by anaphase I the Mtw1^MIS12^-tdTomato signal was undetectable in *ctf19Δ cse4^CENP-A^-S22, S40D* cells (Figure 5B-C). Additionally, Cse4^CENP-A^ phosphomimetics combined with *dsn1-S240, S250D* prevented cells from entering meiosis (Figure S4A). Therefore, the Cse4^CENP-A^ phosphomimetic mutants that rescue kinetochore assembly in the absence of Ipl1^Aurora^ ^B^ instead exacerbate the kinetochore defect of *ctf19Δ* cells.

**Figure 5.**
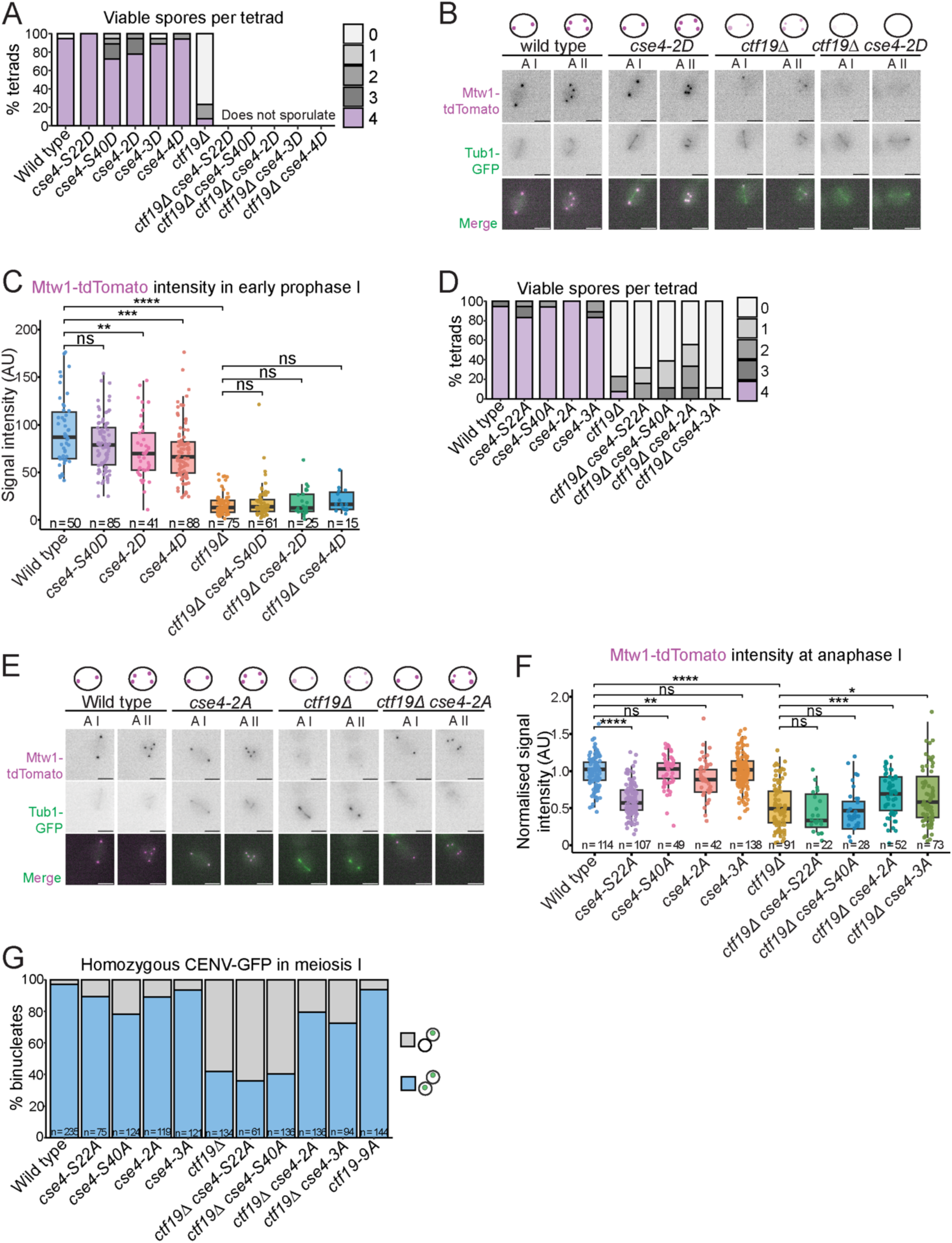
Cse4 phosphorylation precludes kinetochore assembly in *ctf19Δ* cells. (A) Phosphomimetic mutations of *Cse4* prevent sporulation in *ctf19Δ* cells. The number of viable progeny was scored 48 h after tetrad dissection. n = 18 tetrads. (B) Representative images, and illustrative schematics above, from live imaging of Mtw1-tdTomato, Zip1-GFP, and Tub1-GFP at anaphase I and anaphase II. All scale bars represent 2 μm. (C) Phosphomimetic mutations of *Cse4* do not improve kinetochore assembly in *ctf19Δ* cells. Quantification of Mtw1-tdTomato signal intensity from (B) in early prophase I. Meiotic stage determined by Zip1-GFP appearance. All scale bars represent 2 μm. In (C) and (F), boxplots represent interquartile ranges (IQR), and the middle line is the median. Whiskers extend to the furthest data points within 1.5 IQR from the 25th and 75th percentiles; data points outside this range are plotted individually as outliers; Mann-Whitney test. (D) Phosphonull mutations of *Cse4* mildly improve viability of *ctf19Δ* cells. The number of viable progeny was scored 48 h after tetrad dissection. n = 18 tetrads. (E) Representative images, and illustrative schematics above, from live imaging of Mtw1-tdTomato, Zip1-GFP, and Tub1-GFP at anaphase I and anaphase II. All scale bars represent 2 μm. (F) *cse4-2A* rescues the kinetochore assembly defect of *ctf19Δ* cells during the meiotic divisions. Quantification of Mtw1-tdTomato signal intensity from (E) at anaphase I. Meiotic stage determined by Tub1-GFP morphology. (G) *cse4-2A* rescues the meiosis I segregation defect of *ctf19Δ* cells. The percentage of binucleate cells with both copies of chromosome V marked with GFP at the *URA3* locus with the indicated GFP dot segregation was determined. n > 61 binucleates for each strain.

The detrimental effect of Cse4^CENP-A^ phosphomimetic mutants in *ctf19Δ* cells suggested that the Cse4^CENP-A^ NTD may need to remain unphosphorylated to allow kinetochore assembly in *ctf19Δ* cells. Consistently, we found that *cse4^CENP-A^-S22A* or *cse4^CENP-A^-S40A* mildly improved sporulation efficiency and spore viability in *ctf19Δ*, while the combined *cse4^CENP-A^-2A* mutant led to a greater improvement (Figure 5D; S4B). To test whether enhanced kinetochore assembly underlies this rescue, we live imaged Mtw1^MIS12^-tdTomato. Strikingly, the *cse4^CENP-A^-2A* phosphonull mutations significantly increased Mtw1^MIS12^-tdTomato signal intensity in *ctf19Δ* cells, but only during anaphase I and anaphase II, with no rescue observed at meiotic entry or prophase exit (Figure 5E-F; S4C-D). The rescue observed at anaphase I by *cse4^CENP-A^-2A* was abrogated when combined with *dsn1-S240, S250D,* further supporting that Ipl1^Aurora^ ^B^ phosphorylation of its targets is detrimental in the absence of Ctf19^CENP-P^ (Figure S4E-F). Moreover, segregation of homozygous GFP-labelled chromosome V in meiosis I was improved from 41% in *ctf19Δ* to 79% in *ctf19*^CENP-P^*Δ cse4^CENP-A^-2A* (Figure 5G). Despite rescuing kinetochore assembly and meiosis I chromosome segregation, *cse4^CENP-A^-2A* only modestly improved the spore viability of *ctf19Δ* cells (Figure 5D). This is likely explained by the additional role of Ctf19^CENP-P^ in centromeric cohesin loading, which is dispensable for meiosis I segregation but required for accurate segregation during meiosis II (Hinshaw *et al*, 2017; Borek *et al*, 2021). Accordingly, *cse4^CENP-A^-2A* improved segregation in *ctf19Δ* cells almost to the level of *ctf19-9A,* which is specifically defective in targeting cohesin to centromeres, while retaining functional kinetochores (Figure 5G). We conclude that Cse4^CENP-A^ NTD domain phosphorylation at residues S22 and S40 precludes kinetochore assembly in *ctf19Δ* cells. This provides further evidence that Ctf19^CENP-P^ and Ipl1^Aurora^ ^B^ function in separate non-redundant meiotic kinetochore assembly pathways. Mimicking constitutive phosphorylation of Ipl1^Aurora^ ^B^ targets restores kinetochore integrity in the absence of Ipl1^AuroraB^, but is catastrophic in the absence of Ctf19^CENP-P^. Conversely, preventing phosphorylation of Ipl1^Aurora^ ^B^ sites in Cse4^CENP-A^ rescues the kinetochore defect of *ctf19^CENP-P^Δ* cells.

### An alternative Okp1^CENP-Q^-Ame1^CENP-U^ kinetochore assembly pathway is essential after exit from meiotic prophase

Why does Cse4 NTD phosphorylation prevent kinetochore assembly in *ctf19Δ* cells? In addition to binding Scm3 in a phosphorylation-dependent manner, Cse4 END also binds directly to Okp1^CENP-Q^-Ame1^CENP-U^ (Anedchenko *et al*, 2019; Hinshaw & Harrison, 2019; Dendooven *et al*, 2023; Deng *et al*, 2023; Popchock *et al*, 2025). Although this interaction does not occur in the context of the fully assembled Ctf19c^CCAN^, substantial evidence suggests that Okp1^CENP-Q^-Ame1^CENP-U^ can promote kinetochore assembly through this alternative pathway that is independent of the other Ctf19c^CCAN^ components. In mitotic cells, this alternative mode of kinetochore assembly becomes essential only in the absence of the non-essential components of the Ctf19c^CCAN^ (Samel *et al*, 2012). We tested the effect of point mutations in the interface between Cse4^CENP-A^ and Ame1^CENP-U^ in meiosis. *cse4^CENP-A^-L41A, cse4^CENP-A^-R37A,* and *ame1^CENP-U^-I195Y* had an impaired spore viability although markedly less severe than *ctf19Δ* cells (Figure S5A). Strikingly, Mtw1^MIS12^-tdTomato intensity was significantly reduced in *cse4^CENP-A^-L41A, cse4^CENP-A^-R37A,* and *ame1^CENP-U^-I195Y* compared to wild type from early prophase I through to anaphase II (Figure 6A-B). However, in direct contrast to *ctf19Δ* cells, no kinetochore defect was observed at meiotic entry (Figure S5B-C). Therefore, direct Okp1^CENP-Q^-Ame1^CENP-U^ binding to Cse4^CENP-A^, a configuration that is independent of Okp1^CENP-Q^-Ame1^CENP-U^’s role in the context of fully assembled Ctf19c^CCAN^, has a role during meiosis after prophase entry, but is dispensable at meiotic entry.

**Figure 6.**
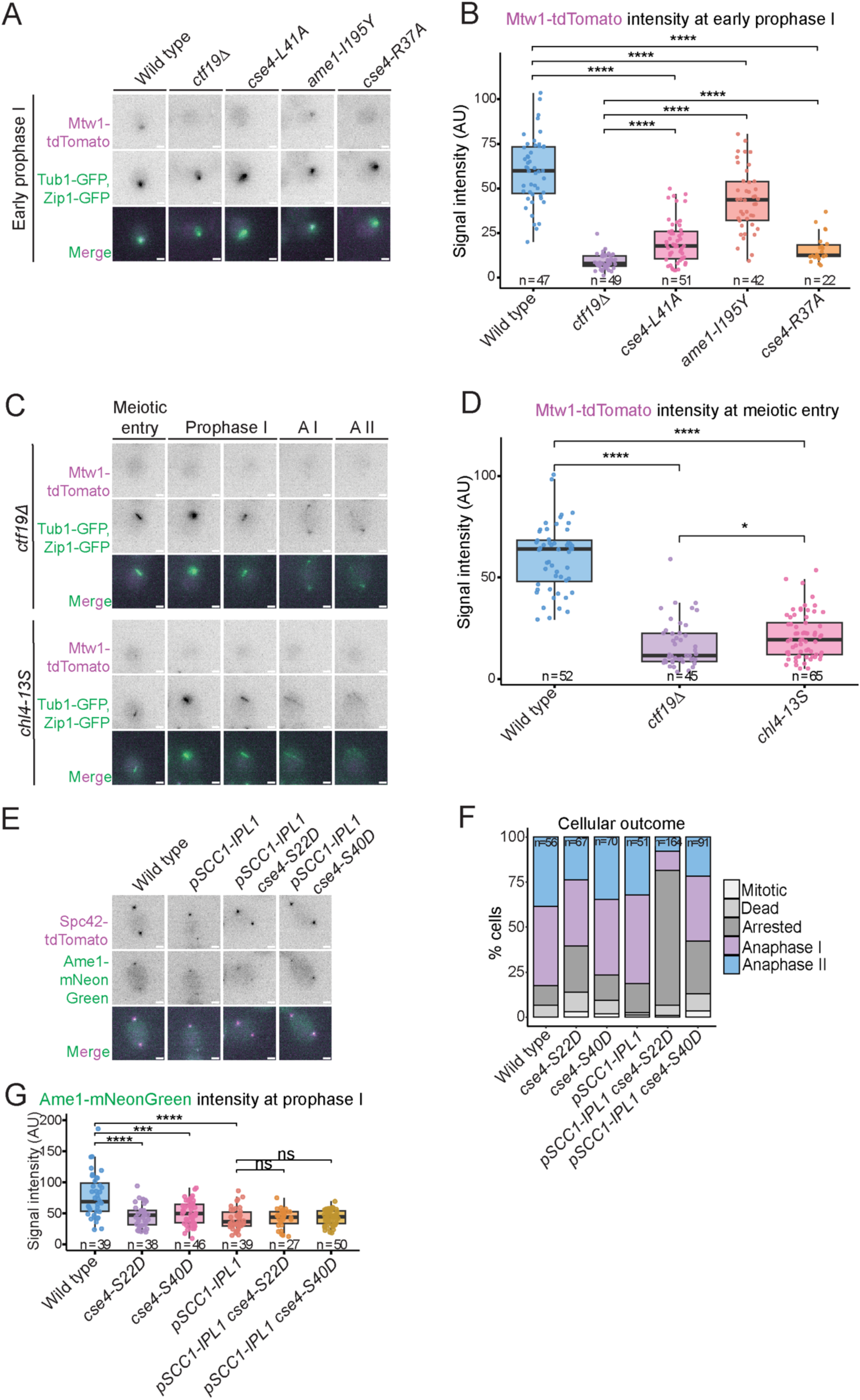
A fully assembled Ctf19c^CCAN^ is required at meiotic entry, while a Ctf19c^CCAN^-independent Okp1^CENP-Q^-Ame1^CENP-U^ kinetochore assembly pathway becomes essential in prophase I to maintain kinetochore integrity. (A) Representative images from live imaging of Mtw1-tdTomato, Tub1-GFP, and Zip1-GFP at early prophase I. Meiotic stage determined by Zip1-GFP appearance. All scale bars represent 2 μm. (B) Mutations preventing the Cse4^CENP-A^-Okp1^CENP-Q^-Ame1^CENP-U^ interaction disrupt kinetochore assembly from prophase I. Quantification of Mtw1-tdTomato signal intensity from (A) at early prophase I. (C) Representative images from live imaging of Mtw1-tdTomato, Tub1-GFP, and Zip1-GFP. All scale bars represent 2 μm. (D) Mutations preventing the interaction of Chl4^CENP-N^ with nucleosomal DNA disrupt kinetochore assembly from meiotic entry. Quantification of Mtw1-tdTomato signal intensity from (C) at meiotic entry. (E) Representative images from live imaging of Spc42-tdTomato and Ame1-mNeonGreen at anaphase I. All scale bars represent 2 μm. (F) Tagging Ame1 prevents meiotic progression in *pSCC1-IPL1* cells with *cse4* phosphomimetic mutations. Percentage of cells reaching each outcome as indicated. (G) Quantification of Ame1-mNeonGreen signal intensity from (E) at prophase I. Meiotic stage determined by Spc42-tdTomato morphology. In (B), (D), and (G), boxplots represent interquartile ranges (IQR), and the middle line is the median. Whiskers extend to the furthest data points within 1.5 IQR from the 25th and 75th percentiles; data points outside this range are plotted individually as outliers; Mann-Whitney test.

### Fully assembled Ctf19c^CCAN^ is critical for kinetochore integrity at meiotic entry

Cells lacking Ctf19^CENP-P^, or other non-essential Ctf19c^CCAN^ components, lose kinetochore integrity at meiotic entry (this study and (Borek *et al*, 2021)), while Ctf19^CENP-P^ becomes dispensable for kinetochore assembly at prophase exit. Instead, an alternative kinetochore assembly pathway involving direct interaction between Okp1^CENP-Q^-Ame1^CENP-U^ is critical at prophase exit. Together, these observations suggest that, at meiotic entry, kinetochore assembly via direct binding of Okp1^CENP-Q^- Ame1^CENP-U^ to Cse4 may not be available. This predicts that, at meiotic entry, kinetochore assembly will be dependent on the fully assembled Ctf19c^CCAN^. In the context of fully assembled Ctf19c^CCAN^, Cse4 binds to Chl4^CENP-N^, and mutation of 13 Arg and Lys residues in a Chl4^CENP-N^ binding groove that interact with one of the unwrapped DNA ends emanating from the Cse4 nucleosome (*chl4^CENP-N^-13S*) can disrupt this interaction (Yan *et al*, 2019). *chl4^CENP-N^-13S* cells are viable but show synthetic sickness with *cse4^CENP-A^-R37A,* which disrupts Okp1^CENP-Q^-Ame1^CENP-U^ binding (Yan *et al*, 2019), consistent with redundant roles of the Okp1^CENP-Q^-Ame1^CENP-U^-only kinetochore assembly mode and the fully assembled Ctf19c^CCAN^ in mitosis. To test the importance of fully assembled Ctf19c^CCAN^ in meiosis, we examined Mtw1^MIS12^-tdTomato intensity in *chl4^CENP-N^-13S* cells and found that levels were very low from meiotic entry, similar to *ctf19Δ* cells (Figure 6C-D). We conclude that fully assembled Ctf19c^CCAN^ is essential to maintain kinetochore integrity at meiotic entry and that the alternative Okp1^CENP-Q^-Ame1^CENP-U^ pathway is not functional at this time.

Furthermore, we tested whether the Cse4^CENP-A^ phosphomimic mutants rescue Ame1^CENP-U^ at meiotic kinetochores in the absence of Ipl1^Aurora^ ^B^. Ame1^CENP-U^-mNeonGreen was live imaged and found to decrease in intensity at *pSCC1-IPL1^Aurora B^* kinetochores (Figure 6E-G). While 82% of wild type and 81% of *pSCC1-IPL1^Aurora^ ^B^* cells completed at least anaphase I, this was mildly decreased in *pSCC1-IPL1 cse4^CENP-A^-S40D* cells to 58% and strongly decreased in *pSCC1-IPL1^Aurora^ ^B^ cse4^CENP-A^-S22D* cells to only 18% (Figure 6F). Accordingly, neither *cse4^CENP-A^-S22D* nor *cse4^CENP-A^-S40D* rescued the loss of Ame1^CENP-U^-mNeonGreen despite the strong rescue of Mtw1^MIS12^-tdTomato in meiosis (Figure 6G; Figure 4D-F). This discrepancy may be attributed to the mNeonGreen tag interfering with Ame1^CENP-U^ function. Interestingly, however, *cse4^CENP-A^-S40D* and *pSCC1-IPL1 cse4^CENP-A^-S40D* cells were not as sensitive to Ame1^CENP-U^-mNeonGreen, suggesting different dependencies on Ame1^CENP-U^. While the reduced function of Ame1^CENP-U^-mNeonGreen may be tolerated in the wild type, the *cse4^CENP-A^-S22D* cells may depend more strongly on Ame1^CENP-U^ to build a kinetochore, compared to *cse4^CENP-A^-S40D* cells. We envisage this to be due to a stronger reliance on Ame1^CENP-U^ function within the Ctf19c^CCAN^, due to Cse4^CENP-A^ NTD phosphorylation diminishing the direct recruitment of Okp1^CENP-Q^- Ame1^CENP-U^. This requirement becomes accentuated in the absence of Ipl1^Aurora^ ^B^ activity, as unphosphorylated Dsn1^DSN1^ would cause Mtw1^MIS12^c autoinhibition, preventing Mif2^CENP-C^ from recruiting Mtw1^MIS12^c.

While *cse4^CENP-A^-2A* rescued Mtw1^MIS12^-tdTomato signal in *ctf19Δ* cells (Figure 5F), it did not rescue Ame1^CENP-U^-mNeonGreen (Figure S5E-F). Again, C-terminal tagging of Ame1^CENP-U^ may be interfering with the rescue. We observed 70% of wild type and 75% of *cse4^CENP-A^-2A* cells reaching at least anaphase I, compared to 10% in *ctf19Δ* cells, however, this was only mildly increased to 23% in *ctf19Δ cse4^CENP-A^-2A* cells carrying Ame1^CENP-U^-mNeonGreen (Figure S5D-F), while cells lacking this tag achieved 76 % sporulation efficiency (Figure S4B). This suggests that *ctf19Δ cse4^CENP-A^-2A* cells are highly dependent on Ame1^CENP-U^ function, and that mild disruption of Ame1^CENP-U^ function can be tolerated in wild-type cells but aggravates the kinetochore assembly defect in *ctf19Δ* cells.

### Kinetochores are dependent on three MIND recruitment pathways to maintain integrity throughout meiosis

In addition to Ame1^CENP-U^, Mif2^CENP-C^ can also recruit the Mtw1^MIS12^ complex via an interaction requiring its N-terminus (Hornung *et al*, 2014; Dimitrova *et al*, 2016; Killinger *et al*, 2020). We tested whether the Mif2^CENP-C^ connection to the outer kinetochore is essential in meiosis despite being dispensable in mitosis.

We found that an N-terminal truncation mutant, *mif2ΔN,* had severely impaired spore viability, with only 11% of tetrads producing spores, albeit at reduced number (Figure 7A). Mutation of a conserved arginine reported to disrupt binding to Mtw1^MIS12^, *mif2-R11A,* mildly impaired spore viability (Figure 7A). To test whether defective kinetochore assembly is responsible for the impaired spore viability, Mtw1^MIS12^-tdTomato was live imaged. The Mtw1^MIS12^-tdTomato signal intensity was significantly decreased in *mif2ΔN* and *mif2-R11A* compared to wild type from meiotic entry and remained low throughout meiosis (Figure 7B-C). Therefore, the Mif2^CENP-C^ connection to the Mtw1^MIS12^ complex is essential from meiotic entry.

**Figure 7.**
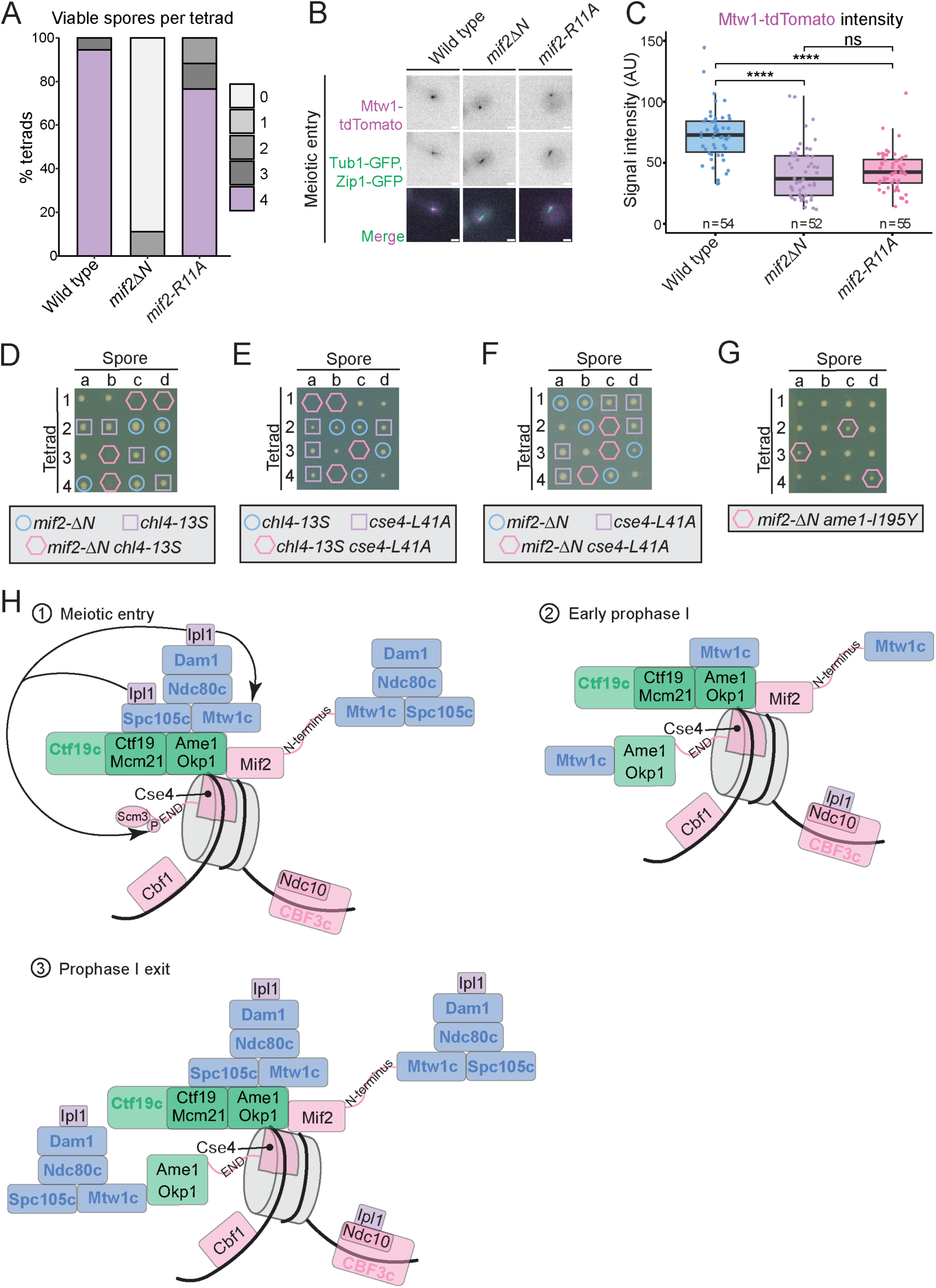
Kinetochores are dependent on three MIND recruitment pathways to maintain integrity throughout meiosis. (A) Preventing Mif2 interaction with Mtw1c impairs spore viability. The number of viable progeny was scored 48 h after tetrad dissection. n = 18 tetrads. (B) Representative images from live imaging of Mtw1-tdTomato, Tub1-GFP, and Zip1-GFP. All scale bars represent 2 μm. (C) Preventing Mif2 interaction with Mtw1c disrupts kinetochore assembly at meiotic entry. Quantification of Mtw1-tdTomato signal intensity from (B) at meiotic entry. Boxplots represent interquartile ranges (IQR), and the middle line is the median. Whiskers extend to the furthest data points within 1.5 IQR from the 25th and 75th percentiles; data points outside this range are plotted individually as outliers; Mann-Whitney test. (D-G) Tetrad dissection and genotype analysis of (D) *mif2ΔN* and *chl4-13S*; (E) *chl4-13S* and *cse4-L41A*; (F) *mif2ΔN* and *cse4-L41A*; (G) *mif2ΔN* and *ame1-I195Y*. (H) Model of kinetochore assembly pathways in meiosis. (1) At meiotic entry, fully assembled Ctf19c^CCAN^ and Mif2 recruit the outer kinetochore. Ipl1 localisation at the Dam1c and Spc105c allows phosphorylation of Dsn1 and Cse4, the latter favouring Scm3 interaction with Cse4 END. (2) In early prophase I, outer kinetochore shedding leads to a relative increase in the Ipl1 pool at Ndc10. Cse4 dephosphorylation promotes Okp1^CENP-Q^-Ame1^CENP-U^ binding at the END in lieu of Scm3. (3) By prophase I exit, the outer kinetochore is reassembled, and Ipl1 localises to Ndc10 and Dam1c. All three pathways recruiting Mtw1c are active.

We have shown that the Ctf19c^CCAN^- and Mif2^CENP-C^-dependent kinetochore integrity pathways are required from meiotic entry, while the Cse4^CENP-A^-Okp1^CENP-Q^- Ame1^CENP-U^ pathway is required from early prophase I. Given that the individual pathways are dispensable in mitosis, we tested whether they are redundant. We used *chl4-13S* to disrupt the Ctf19c^CCAN^ pathway, *mif2DN* to disrupt the Mif2^CENP-C^ pathway, and either *cse4-L41A* or *ame1-I195Y* to disrupt the Cse4^CENP-A^-Okp1^CENP-Q^- Ame1^CENP-U^ pathway. Haploid cells with these single mutants were all viable, as expected. To assess the viability of double mutant combinations, we crossed the corresponding haploids, induced meiosis in the resultant heterozygous diploids and genotyped the haploid spores. No viable double mutant spores were recovered for *chl4-13S mif2ΔN* (Figure 7D), *chl4-13S cse4-L41A* (Figure 7E) or *mif2ΔN cse4-L41A* (Figure 7F), revealing synthetic lethality between these mutations. However, *mif2ΔN ame1-I195Y* spores were viable but formed smaller colonies than either single mutant (Figure 7G). This was consistent with the milder meiotic kinetochore assembly defect of *ame1-I195Y* compared to *cse4-L41A* (Figure 6F), further suggesting this point mutation does not fully disrupt the Cse4^CENP-A^ END interaction with Okp1^CENP-Q^- Ame1^CENP-U^. Together, these findings indicate that at least two connections to Mtw1^MIS12^c are required for viability in mitosis, whereas all three are necessary to maintain kinetochore integrity throughout meiosis.

## Discussion

### Ctf19c^CCAN^ and Ipl1^AuroraB^ independently maintain meiotic kinetochore integrity

Kinetochores take on additional functions in meiosis (Koch & Marston, 2025). We previously reported that meiotic kinetochore integrity is highly sensitive to the loss of Ctf19^CENP-P^, in contrast to mitotic cells where Ctf19^CENP-P^ is not essential (Borek *et al*, 2021). Here, we identify three pathways connecting the centromere to the outer kinetochore: one through the assembled Ctf19c^CCAN^, a second through direct binding of Okp1^CENP-Q^-Ame1^CENP-U^ to Cse4^CENP-A^, and a third through Mif2^CENP-C^ (Figure 7H). Ctf19^CENP-P^ is required from meiotic entry until prophase I exit, after which kinetochore integrity and sporulation can occur in its absence. Mutations in non-essential Ctf19c^CCAN^ components are synthetic lethal with mutations in CPC components (Makrantoni *et al*, 2017; Campbell & Desai, 2013; Ng *et al*, 2009). This genetic interaction has been attributed to recruitment of Ipl1^Aurora^ ^B^ by Ctf19^CENP-P^ through Sli15^INCENP^ in mitosis (Fischböck-Halwachs *et al*, 2019; García-Rodríguez *et al*, 2019). However, we find no evidence for a Ctf19^CENP-P^-dependent pool of Ipl1^Aurora^ ^B^ in meiosis, despite Ipl1^Aurora^ ^B^ being required throughout meiosis for kinetochore assembly. Instead, at meiotic entry and prophase I exit, Ipl1^Aurora^ ^B^ localises near Dam1c and Spc105^KNL1^c at the outer kinetochore, consistent with mitotic studies (Dudziak *et al*, 2025).

We therefore propose that Ctf19^CENP-P^ and Ipl1^Aurora^ ^B^ function in distinct pathways to maintain kinetochore integrity. Our findings also support a model in which dynamic phosphorylation of Ipl1^Aurora^ ^B^ target sites regulates kinetochore assembly. Cse4^CENP-A^ and Dsn1^DSN1^ phosphorylation promotes kinetochore assembly in early meiosis, but constitutive phosphorylation is detrimental in *ctf19Δ* cells, perhaps because Cse4^CENP-A^ NTD phosphorylation impedes Okp1^CENP-Q^-Ame1^CENP-U^ binding. Despite this, phosphomimetic Cse4^CENP-A^ mutants are viable, suggesting that they weaken, rather than completely abolish, Okp1^CENP-Q^-Ame1^CENP-U^ binding.

### Meiosis requires all three centromere-outer kinetochore connections at different stages

Overall, our data support a model in which the relative contributions of the three connections change throughout meiosis (Figure 7H). At meiotic entry, both the Ctf19^CENP-P^-dependent and Mif2^CENP-C^-dependent pathways are required for Mtw1c^MIS12c^ recruitment. Cse4^CENP-A^ interaction with the Chl4^CENP-N^ DNA-binding groove is also essential, indicating a requirement for proper Ctf19c^CCAN^ assembly. At this stage, outer kinetochore-associated CPC may phosphorylate the Cse4^CENP-A^ NTD and Dsn1^DSN1^, stabilising Cse4^CENP-A^ through Scm3^HJURP^ and relieving Mtw1^MIS12^ autoinhibition, respectively. During early prophase I, the outer kinetochore disassembles and its associated CPC is lost, while the CPC pool at Ndc10 relatively increases. We propose that reduced Cse4^CENP-A^ NTD phosphorylation decreases its affinity for Scm3^HJURP^ and favours Okp1^CENP-Q^-Ame1^CENP-U^ binding instead, consistent with the decline in kinetochore-localised Scm3^HJURP^ during prophase I. The rescue of Mtw1^MIS12^ levels in *ctf19Δ cse4-2A* cells from prophase I exit further supports this possibility. However, *cse4-2A* does not restore kinetochore assembly in *ctf19Δ* at meiotic entry, suggesting additional requirements for this connection.

Loss of CPC upon outer kinetochore shedding during prophase I may also lead to Mtw1^MIS12^ autoinhibition, weakening the Mif2^CENP-C^ connection and increasing dependence on Okp1^CENP-Q^-Ame1^CENP-U^. Alternatively, kinetochore remodelling may generally increase the requirement for multiple connections. At prophase I exit, the outer kinetochore and its associated CPC are restored. In addition to facilitating error correction, Ipl1^Aurora^ ^B^ may then relieve Mtw1^MIS12^c autoinhibition, allowing the Mif2^CENP-C^ -dependent connection to function alongside the Cse4^CENP-A^-Okp1^CENP-Q^-Ame1^CENP-U^ pathways. These alternative connections could explain why Ctf19c^CCAN^ becomes dispensable after prophase I exit (Figure 7H).

The changing requirement for these connections may reflect the extensive remodelling of meiotic kinetochores (Koch & Marston, 2025; Borek *et al*, 2021). During premeiotic S phase, kinetochores promote pericentromeric cohesin, which, together with the kinetochore itself, suppresses crossover recombination (Fernius & Marston, 2009; Hinshaw *et al*, 2017; Vincenten *et al*, 2015). During prophase, upon outer kinetochore disassembly, telomeres attach to microtubules to form the so-called prophase bouquet (Meyer *et al*, 2015; Miller *et al*, 2012; Hayashi *et al*, 2006). Ipl1^AuroraB^ promotes this assembly, potentially through the Spc105^KNL1^-associated pool (Meyer *et al*, 2015; Borek *et al*, 2021; Chen *et al*, 2020). At prophase exit, Ndc80^NDC80^ re-expression enables outer kinetochore reassembly (Chen *et al*, 2017), while Cdc5^Plk1^ promotes monopolin recruitment and sister kinetochore co-orientation (Rabitsch *et al*, 2003). These transitions coincide with a shift from low CDK and Cdc5^Plk1^ activity during prophase to increased kinase activity during the meiotic divisions (Koch *et al*, 2024; Carlile & Amon, 2008; Celebic *et al*, 2024). Thus, rather than simply providing redundancy, multiple centromere-outer kinetochore connections may enable kinetochores to accommodate the changing structural and regulatory requirements of meiosis.

### Alternative kinetochore connections explain the essentiality of Okp1^CENP-Q-^Ame1^CENP-U^

Our findings provide an explanation for why Okp1^CENP-Q^ and Ame1^CENP-U^ are the only essential Ctf19c^CCAN^ subunits, while the N-terminal Mtw1c^MIS12c^-binding domain of Mif2^CENP-C^ is dispensable. Okp1^CENP-Q^-Ame1^CENP-U^ contribute to two of the three connections between the centromere and outer kinetochore: one through the assembled Ctf19c^CCAN^ and another through direct binding to the Cse4 ^CENP-A^ END (Anedchenko *et al*, 2019; Hinshaw & Harrison, 2019; Deng *et al*, 2023; Dendooven *et al*, 2023). Loss of Okp1-Ame1 would therefore compromise both connections, leaving only the Mif2^CENPC^-dependent pathway, which is insufficient for viability. Consistent with this, our mitotic genetic analysis indicates that loss of one connection is tolerated, whereas disruption of two connections results in synthetic lethality or severe growth defects.

Importantly, direct Cse4^CENP-A^-END-Okp1^CENP-Q^-Ame1^CENP-U^ binding is required during meiosis even when Ctf19c^CCAN^ assembly is intact, indicating that it is not simply a backup connection. Structural studies have revealed that the configuration of Okp1^CENP-Q^-Ame1^CENP-U^ when bound directly to the Cse4^CENP-A^ END is incompatible with its arrangement within the assembled Ctf19c^CCAN^ (Dendooven *et al*; Deng *et al*, 2023; Hinshaw & Harrison, 2019). This raises the possibility that kinetochores adopt alternative architectures, although whether these configurations coexist within individual kinetochores or are used at different stages remains unknown. Our findings suggest that such alternative connections make distinct contributions to kinetochore integrity during meiosis.

### Evolutionary diversity in centromere-outer kinetochore connections

Although the outer kinetochore machinery is broadly conserved, the connections linking it to centromeric chromatin are remarkably diverse across eukaryotes. In vertebrates, CENP-C and CENP-T provide major pathways for outer kinetochore recruitment (Gascoigne *et al*, 2011; Hori *et al*, 2013), whereas CENP-T is dispensable in budding yeast, where Okp1^CENP-Q^–Ame1^CENP-U^ provides an additional connection (Bock *et al*, 2012; Schleiffer *et al*, 2012). Other organisms have more divergent architectures: *Drosophila* has lost most canonical CCAN components and relies primarily on CENP-C (Przewloka *et al*, 2011), while some lineages lack recognisable CENP-A or CENP-C homologues altogether (van Hooff *et al*, 2017). Interestingly, Okp1^CENP-Q^–Ame1^CENP-U^ does not appear to have been retained independently in organisms that have lost other CCAN components (van Hooff *et al*, 2017). This suggests that even though Okp1^CENP-Q^–Ame1^CENP-U^ can form an independent connection, this works alongside, rather than instead of, the wider CCAN architecture. Thus, although organisms have evolved different mechanisms for connecting centromeric chromatin to the outer kinetochore, these connections may not be freely interchangeable. Our findings highlight how alternative connections can provide flexibility within an organism, while also suggesting constraints on the architectures that can support chromosome segregation.

## Methods

### Yeast strains and plasmids

All yeast strains are SK1 derivatives and are listed in Table S1. Gene deletions, gene tags, and promoter replacements were introduced using standard PCR-based methods (Knop *et al*, 1999; Longtine *et al*, 1998), with the exception of the *CSE4-mNeonGreen* allele, which was generated by CRISPR. Point mutations in *CSE4, CHL4,* and *MIF2,* and the *mif2(Δ2-30)* alleles were also generated by CRISPR (Vinod *et al*, 2026).

### CSE4-mNeonGreen construction

Cse4 was internally tagged with mNeonGreen by inserting the tag flanked by two linkers between L81 and E82. The fluorescent tag was amplified from AMp1604 (pFA6a-mNeonGreen-KILEU2) using primers each with 100 bp homology to *CSE4* (AMo12343, AMo12289). Primers encoding sgRNA (AMo7441, AMo7442) allowing Cas9 to cut at *CSE4* G79 were cloned into AMp1278 (pWS082) to produce AMp1285. The sgRNA fragment was amplified from AMp1295 using primers AMo6663, AMo6664. A fragment containing *CAS9* and an *HPH* marker was amplified from AMp1285 using primers AMo6723, AMo6724. All three fragments were transformed into yeast, and correct integrants were confirmed by sequencing.

### Point mutations in *CSE4, CHL4,* and *MIF2*

Point mutations were generated by CRISPR as described in “*CSE4-mNeonGreen* construction”. Guide RNAs targeting the region of interest were designed on Benchling and cloned into AMp1278 (pWS082) by Golden Gate assembly. Homology-directed repair templates containing the desired mutation and silent mutations to disrupt the PAM site were amplified by PCR. Yeast were transformed with PCR fragments encoding Cas9, the sgRNA, and the repair template. Correct mutant strains were identified by allele-specific PCR and sequencing. Plasmids generated in this study are given in Table S2.

### *mif2(Δ2-30)* construction

The *mif2ΔN* truncation mutant was generated by CRISPR as described above using a homology-directed repair template containing the *MIF2* sequence truncated from amino acid 2-30. Yeast were transformed with PCR fragments encoding Cas9, the sgRNA, and the repair template. Correct mutant strains were identified by allele-specific PCR and sequencing. Plasmids generated in this study are given in Table S2.

## Method Details

### Yeast growth conditions

#### Meiotic induction

Yeast strains were recovered from −70 °C and grown for 16 h on YPG plates (1 % yeast extract, 2 % bactopeptone, 2.5 % glycerol, and 2 % agar), and then for 8 h on YPD 4 % plates (1 % yeast extract, 2 % bactopeptone, 4% glucose, and 2 % agar). Cells were grown in YPDA (1 % yeast extract, 2 % bactopeptone, 2 % glucose, and 0.3 mM adenine) overnight for 12 h before inoculation at OD_600_ = 0.3 into BYTA (1 % yeast extract, 2 % bactotryptone, 1 % potassium acetate, and 50 mM potassium phthalate) and grown overnight to reach an OD_600_ ≥ 5. Cells were harvested, washed twice with water, and resuspended into SPO (0.3 % potassium acetate, pH 7.0) at OD_600_ = 2.3. SPO cultures were shaken vigorously. All growth steps were carried out at 30 °C. For synchronisation at meiotic entry, cells carrying the *pCUP-IME1 pCUP-IME4* alleles were grown for 2 h before release by addition of 25 μM copper (II) sulphate, as previously described (Berchowitz *et al*, 2013). For synchronisation during the meiotic divisions, cells carrying *pGAL-NDT80* were grown for 6 h before release from prophase I arrest by addition of 1 μM β-estradiol.

#### Growing strains for TurboID

Yeast strains containing miniTurboID-tagged proteins were induced to undergo meiosis. 50 μM biotin was added to SPO cultures 50 min prior to the time of harvest for efficient labelling.

#### Flow cytometry

150 µL of meiotic culture was fixed in 350 µL 100 % EtOH and stored at 4 °C. Cells were pelleted and resuspended in 1 mL 50 mM Tris-HCl, pH 7.5. Samples were sonicated in the Biorupter (Diagenode) on LOW with 1 cycle of 30 sec ON/30 sec OFF. Cells were incubated in 475 µL 50 mM Tris-HCl, pH 7.5 and 25 µL 20 mg/mL RNaseA at 37 °C overnight. Cells were washed in 1 mL 50 mM Tris-HCl, pH 7.5, resuspended in 500 µL 50 mM Tris-HCl, pH 7.5 with 10 µL 20 mg/mL Proteinase K (Amresco), and incubated at 50 °C for 2 h. Cells were washed with 1 mL 50 mM sodium citrate and stained with 500 µL sodium citrate and 9.17 µL 1 mg/mL propidium iodide in the dark at 4 °C. Samples were sonicated in the Biorupter (Diagenode) immediately before running on flow cytometer on LOW setting with 10 cycles 30 sec ON/30 sec OFF. Samples were analysed using a BD FACS Calibur instrument.

### Microscopy

#### Live cell imaging

Concanavalin A was spread in the wells of an 8-well Ibidi dish and incubated at 30 °C for 15 min. Wells were washed three times. Meiotic SPO cultures were added to the wells, and cells were left to attach by incubation for 20 min at 30 °C. The dish was washed twice, and 400 μL SPO media was added for imaging.

A Zeiss Axio Observer Z1 (Zeiss UK, Cambridge) with a Prior motorised stage was used for live imaging with a Hamamatsu Flash 4 sCMOS camera and Zen 2.3 acquisition software. 2×2 binning was used. Images were acquired every 8-15 min for 12 h. 9 slices were imaged in the z-plane with 0.7 mm spacing. Exposure duration and percentage of light transmitted were optimised for each fluorophore and according to experimental duration.

#### Chromosome segregation assay

Diploid strains with both copies of chromosome V marked with GFP were induced to enter meiosis as described in “Meiotic induction”, and cells were transferred to an 8-well Ibidi glass-bottom dish after ∼4 hours of growth as described in “Imaging chamber preparation”. Cells were live imaged for 8-10 hours, and GFP dots were scored in at least 100 tetranucleate cells for each strain.

#### Fixed cell imaging

100 μL of meiotic culture was fixed in 10 μL 37 % v/v formaldehyde for 8 min at RT. Cells were pelleted and washed in 1 mL 80% ethanol. Cells were pelleted and resuspended in 30 μL 1 μg/mL DAPI or 30 μL PBS and stored at 4 °C until imaging. For imaging, sample tubes were briefly vortexed and 4 μL of cells were placed on a slide that was imaged using a Zeiss Axioplan 2 microscope with 100X Plan ApoChromat NA 1.4 oil lens and Zeiss Immersol oil 518F. 1×1 binning was used. 9 slices were imaged in the z-plane with 0.7 mm spacing.

#### Spindle immunofluorescence

300 μL meiotic culture was spun down, and the pellet resuspended in 500 μL 3.7 % v/v formaldehyde in 0.1 M potassium phosphate at pH 6.4 to fix overnight at 4 °C. Cells were washed 3 times with 1 mL 0.1 M potassium phosphate at pH 6.4 and resuspended in 1 mL 1.2 M sorbitol citrate. Cells were resuspended in digestion mix (200 μL 1.2 M sorbitol citrate, 20 μL glusulase (Perkin Elmer) and 6 μL 10 mg/mL zymolyase (AMS Biotechnology)) and incubated at 30 °C for 1 h or until digestion was complete. Digested cells were washed with 1 mL 1.2 M sorbitol-citrate and resuspended in 30 μL 1.2 M sorbitol citrate. Cells were attached to multi-well polylysine-treated slides and fixed in MeOH for 3 min and in acetone for 10 s, then dried. Rat anti-tubulin primary antibody (Bio-Rad) at 1:50 dilution in PBS/BSA (1 % w/v BSA, 0.04 M K_2_HPO_4_, 0.01 M KH_2_PO_4_) was added to each well and incubated for 1 h at 30 °C. Wells were washed 5 times with PBS/BSA and donkey anti-rat-FITC secondary antibody (Jackson ImmunoResearch) at 1:100 dilution in PBS/BSA was added to each well and incubated for 1 h at 30 °C. Wells were washed a further 5 times with PBS/BSA and 3 μL DAPI-MOUNT (9 mM p-phenylenediamine, 0.04M K_2_HPO_4_, 0.01 M KH_2_PO_4_, 0.15 M NaCl, 0.1 % w/v NaN_3_, 50 ng/mL DAPI, 90 % v/v glycerol) was added to each well and a coverslip placed on the slide before imaging.

#### Sporulation efficiency assay

Diploids were induced to enter meiosis as described in “Meiotic induction”. Sporulation efficiency was determined by light microscopy after 24 hours in SPO medium.

#### Spore viability assay

Viability of meiotic cells was determined by dissecting 18 tetrads of a homozygous diploid carrying the mutation of interest. Viability was scored after 48 hours of recovery at 30 °C.

#### Western blotting

Samples were fixed in trichloroacetic acid for 10 min on ice, washed in acetone, and whole cell extracts were prepared by bead beating in TE-containing protease inhibitors. Proteins were transferred to nitrocellulose membrane as previously described (Barton *et al*, 2022). Antibodies used were rabbit anti-PGK1 (lab stock, 1:50,000), rabbit anti-KAR2 (lab stock, 1:5000), mouse anti-V5 (Bio-Rad, 1:2000), sheep anti-mouse-HRP (GE Healthcare, 1:5000), donkey anti-rabbit-HRP (GE Healthcare, 1:10,000).

#### Immunoprecipitation of streptavidin-coated Dynabeads

400 mL meiotic culture was harvested for each strain after addition of 1 mM PMSF and the cell pellet washed twice in 50 mL PBS supplemented with 1 mM PMSF. Cells were snap frozen as dots and stored at −70 °C. Cells were lysed in a SPEX 6875 freezer mill by 8 rounds of 2 min cooling, 2 min grinding, 10 CPS and the grindate stored at −70 °C. For the pulldown, the grindate was thawed and 2.5 g were resuspended in 10 mL RIPA lysis buffer (50 mM Tris-HCl, pH7.5, 150 mM NaCl, 0.1 % w/v SDS, 0.5 % w/v sodium deoxycholate, 1X v/v NP-40) supplemented with 1 mM DTT, 2 mM PMSF, and 1X EDTA-free Roche protease inhibitor tablet. 10 μL BaseMuncher benzonase and 1 mM MgCl_2_ were added, and the grindate paste was incubated for 1.5 h at 4 °C with rotation. 300 μL Dynabead C1 beads (to immunoprecipitated 3 mg protein per strain) were washed three times with 1 mL RIPA buffer for 5 min each on a rotator and magnetic rack. Clarified lysate was obtained by centrifugation at 4,000 rpm for 20 min at 4 °C and protein concentration was determined by Bradford assay. Equilibrated Dynabead C1 beads were added to clarified lysate containing 3 mg protein in a Protein LoBind Eppendorf and incubated for 2.5 h at 4 °C with rotation. Beads were washed 2X with 1 mL RIPA buffer, 1X with 1 mL 1M KCl, and 6X with 1 mL 20 mM ammonium bicarbonate for 8 min each on a rotator. Proteins were eluted by incubating the beads with 55 μL elution buffer (2X NuPAGE LDS sample buffer, 1X NuPAGE reducing agent, 0.25 mM biotin in SPO, 20 mM DTT) and boiling for 5 min at 95 °C. Eluate was stored at −20 °C for mass spectrometry analysis.

#### Mass spectrometry

Protein samples were briefly separated on an SDS-PAGE gel (NuPAGE Novex 4%– 12% Bis-Tris gel, ThermoFisher, UK), in NuPAGE buffer (MES) and visualized using InstantBlue stain (Sigma Aldrich, UK). The stain was removed by washing 3x with water. Protein bands were excised and gel pieces were destained with 50 mM ammonium bicarbonate (Sigma Aldrich, UK) and 100% v/v acetonitrile (Sigma Aldrich, UK) for 30 min at 37 °C. Gel pieces were reduced in 10 mM dithiothreitol (Sigma Aldrich, UK) for 30 min at 37 °C and alkylated in 55 mM iodoacetamide (Sigma Aldrich, UK) for 20 min at room temperature in the dark. They were then digested overnight at 37 C with 12.5 ng/mL trypsin (Pierce, UK) as described in (Shevchenko *et al*, 1996). Following digestion, samples were diluted with an equal volume of 0.1% TFA and spun onto StageTips. Remaining peptides were eluted from the gel pieces in 40 μL of 80% acetonitrile in 0.1% TFA and concentrated by vacuum centrifugation (Concentrator 5301, Eppendorf, UK). Peptides were resuspended with 80 % ACN and 0.1 % TFA and passed through the StageTips before storing at −20 °C until analysis by LC-MS/MS (Rappsilber *et al*, 2007).

#### Quantitative analysis of mass spectrometry data

LC-MS analyses were performed on an Orbitrap Exploris™ 480 (Thermo Fisher Scientific, UK) on a Data Independent Acquisition (DIA) mode, coupled on-line to an Ultimate 3000 HPLC (Dionex, Thermo Fisher Scientific, UK). Peptides were separated on a 50 cm (2 µm particle size) EASY-Spray column (Thermo Scientific, UK), which was assembled on an EASY-Spray source (Thermo Scientific, UK) and operated constantly at 55 °C. Mobile phase A consisted of 0.1% formic acid in LC-MS grade water, and mobile phase B consisted of 80% acetonitrile and 0.1% formic acid. Peptides were loaded onto the column at a flow rate of 0.3 μL min^-1^ and eluted at a flow rate of 0.25 μL min^-1^ according to the following gradient: 2 to 40% mobile phase B in 150 min and then to 95% in 11 min. Mobile phase B was retained at 95% for 5 min and returned back to 2% a minute after until the end of the run (190 min).

Survey scans were recorded at 120,000 resolution (scan range 350-1650 m/z) with an ion target of 5.0e6, and injection time of 20ms. MS2 was performed in the Orbitrap at 30,000 resolution with a scan range of 200-2000 m/z, maximum injection time of 55 ms, and AGC target of 3.0E6 ions. We used HCD fragmentation with stepped collision energy of 25.5, 27 and 30 and variable isolation windows throughout the scan range, ranging from 10.5 to 50.5 m/z. Narrow isolation windows (10.5-18.5 m/z) were applied from 400-800 m/z and then gradually increased to 50.5 m/z until the end of the scan range. In total we used 45 isolation windows for the reported scan range. The default charge state was set to 3. Data for both survey and MS/MS scans were acquired in profile mode.

The DIA-NN software platform (Demichev *et al*, 2020) version 1.9.2. was used to process the DIA raw files, and search was conducted against the *Saccharomyces cerevisiae,* proteome strain SK1 (from *Saccharomyces* Genome Database, released in 2019). Precursor ion generation was based on the chosen protein database (automatically generated spectral library) with deep-learning-based spectra, retention time and IMs prediction. Digestion mode was set to specific with trypsin, allowing a maximum of two missed cleavages. Carbamidomethylation of cysteine was set as a fixed modification. Oxidation of methionine and acetylation of the N-terminus were set as variable modifications. The parameters for peptide length range, precursor charge range, precursor m/z range and fragment ion m/z range, as well as other software parameters, were used with their default values. The precursor FDR was set to 1%. Spectral library was created with information from the FASTA database.

#### Quantification and statistical analyses

Statistical analyses were performed on R software and are detailed in each figure legend. In all figures, a single asterisk (*) indicates a p-value of less than or equal to 0.05, two asterisks (**) indicate a p-value of less than or equal to 0.01, three asterisks (***) indicate a p-value of less than or equal to 0.001, and four asterisks (****) indicate a p-value less than or equal to 0.0001.

## Acknowledgements

We are grateful to members of the Marston group for helpful discussions. We thank Dilara Kocakaplan for assistance with the TurboID experiments. We gratefully acknowledge the Wellcome Discovery Research Platform for Hidden Cell Biology Proteomics and Light Microscopy Cores. This work was funded through Wellcome Investigator [220780] and Discovery Awards to AM [319314], a UKRI BBSRC EASTBIO studentship to MH [BB/T00875X/1] and a Wellcome Discovery Research Platform Award [226791].

## Author contributions

Conceptualisation – MH, WB, AM; Data curation – MH, CS; Funding Acquisition – AM; Formal Analysis – MH, CS, DK; Investigation – MH, WB; Software – DK; Supervision – AM; Visualisation – MH; Writing – original draft – MH; Writing – review and editing – MH and AM with input from all authors.

---

**Figure S1.**
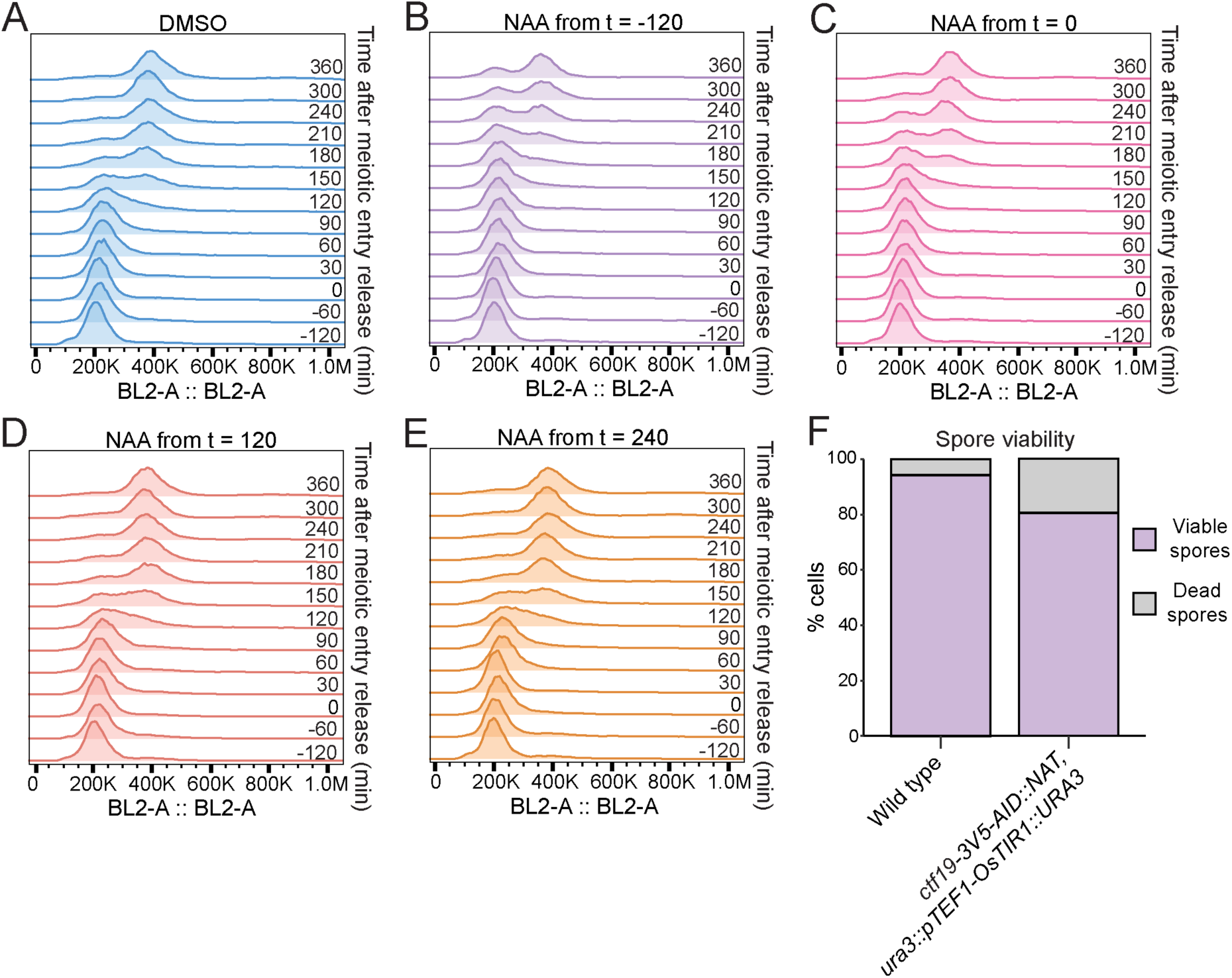
The *ctf19-3V5-AID* allele does not disrupt meiosis progression. Related to Figure 1. (A-E) DNA replication completion is unaffected by Ctf19-3V5-AID depletion. Flow cytometry profiles at each timepoint indicated in relation to meiotic entry release when cells were treated with (A) DMSO; (B) NAA from 2 h before meiotic entry release; (C) NAA from meiotic entry release; (D) NAA from 2 h after meiotic entry release; (E) NAA from 4 h after meiotic entry release. (F) *ctf19-3V5-AID* allele does not strongly impair spore viability. The number of viable progeny was scored 48 h after tetrad dissection. n = 18 tetrads.

**Figure S2.**
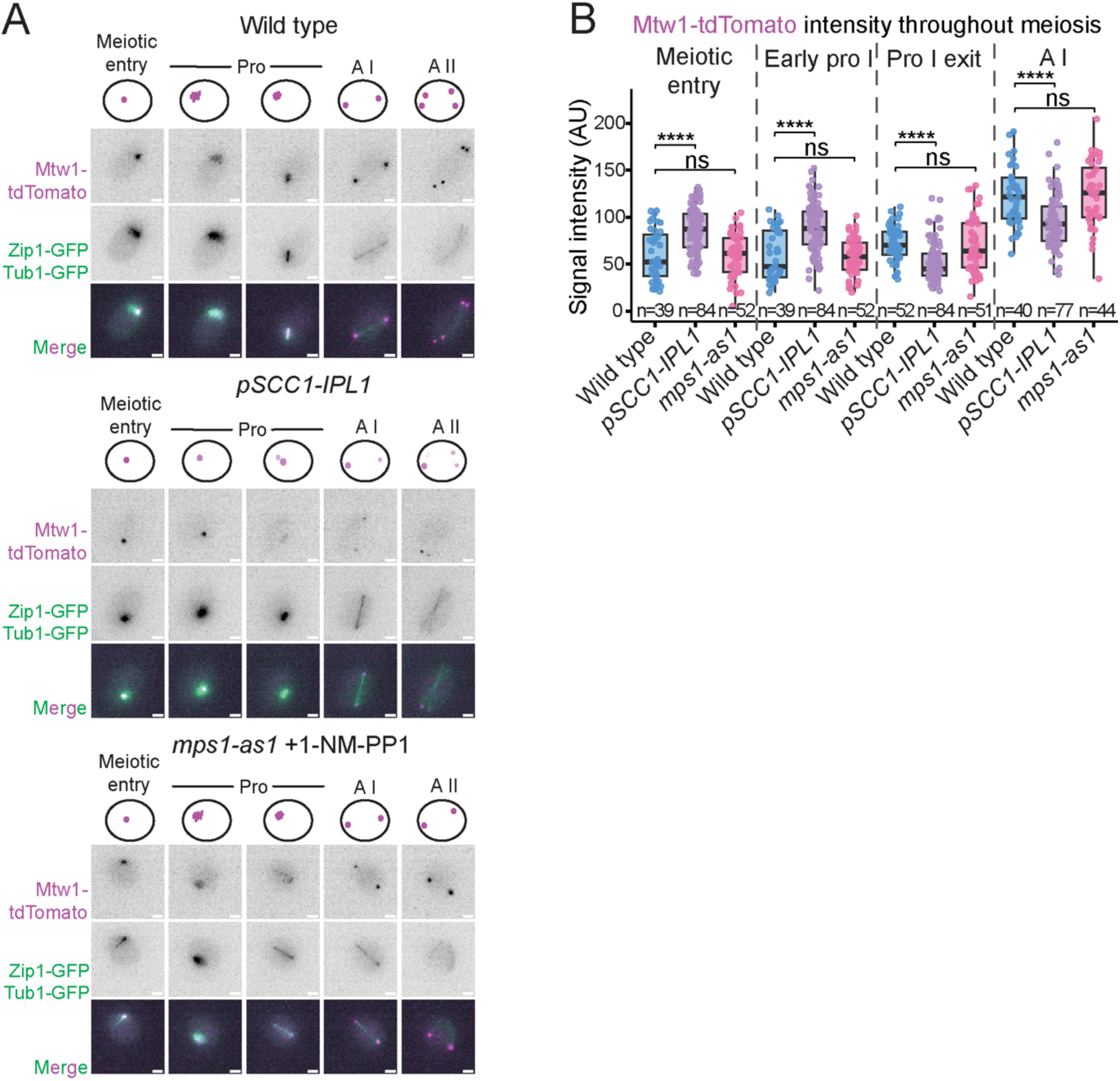
Mps1 kinase activity is not required for meiotic kinetochore integrity. Related to Figure 2. (A) Representative images, and illustrative schematics above, from live imaging of Mtw1-tdTomato, Zip1-GFP, and Tub1-GFP. All scale bars represent 2 μm. (B) Inhibiting Mps1 kinase activity does not cause a kinetochore assembly defect. Quantification of Mtw1-tdTomato signal intensity from (A). Meiotic stage determined by Zip1-GFP appearance and disappearance, and Tub1-GFP morphology. Boxplots represent interquartile ranges (IQR), and the middle line is the median. Whiskers extend to the furthest data points within 1.5 IQR from the 25th and 75th percentiles; data points outside this range are plotted individually as outliers; Mann-Whitney test.

**Figure S3.**
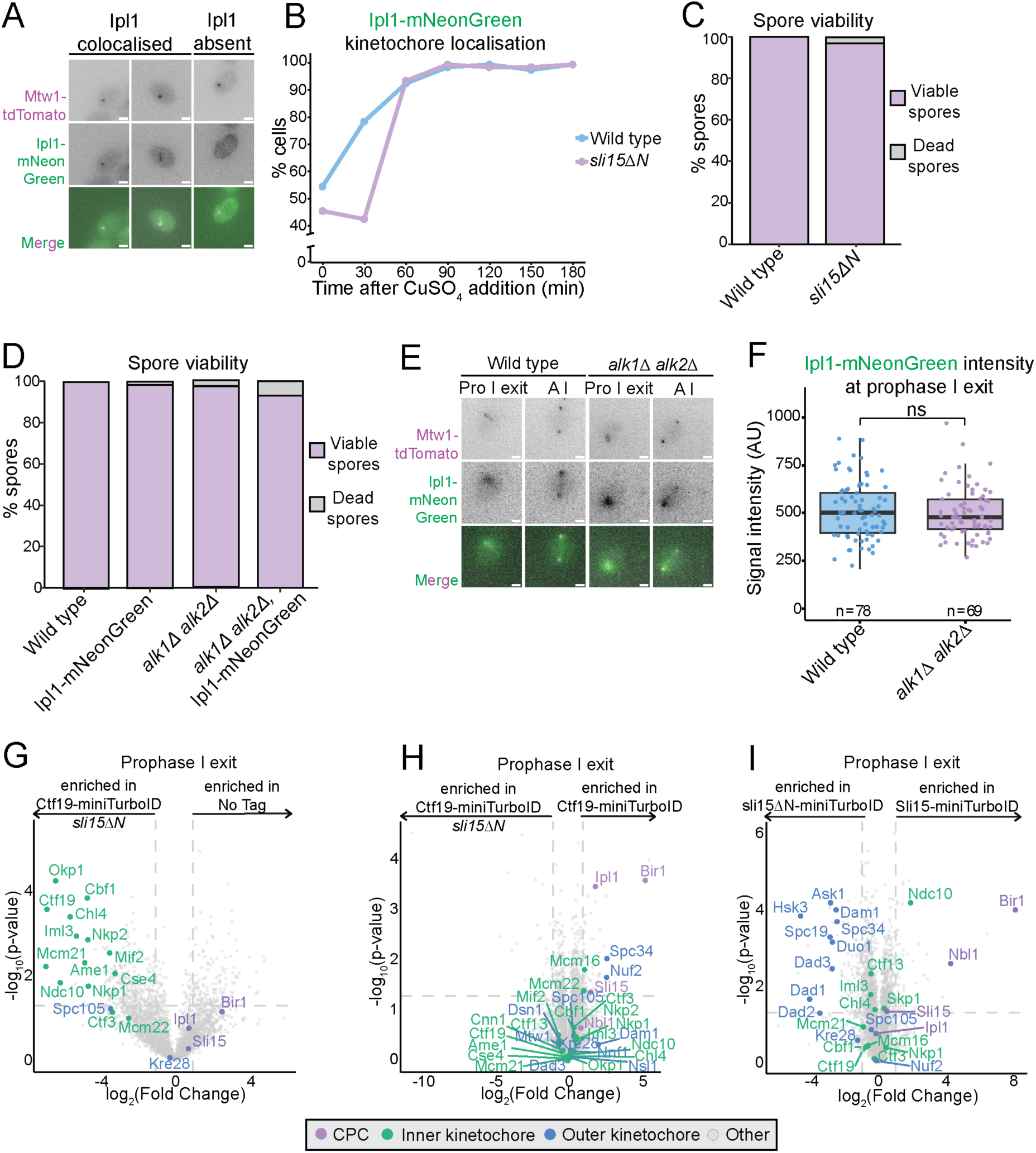
Ctf19 does not mediate CPC recruitment in the absence of Bir1-dependent recruitment pathways in meiosis. Related to Figure 3. (A) Representative fixed cell images of Mtw1-tdTomato and Ipl1-mNeonGreen. All scale bars represent 2 μm. (B) Ipl1 localisation is unperturbed in the absence of Bir1-dependent recruitment pathways. Percentage of cells with Ipl1-mNeonGreen localised with Mtw1-tdTomato from imaging in (A). Diploids containing the *pCUP-IME1 pCUP-IME4* system were induced to undergo meiosis. After 2 h, copper (II) sulfate was added to release cells synchronously into meiotic entry, and samples were taken every 30 min. n = 100 cells per timepoint. (C) *sli15ΔN* does not impair spore viability. The number of viable progeny was scored 48 h after tetrad dissection. n = 18 tetrads. (D) *Ipl1-mNeonGreen* and *alk1Δ alk2Δ* do not impair spore viability. The number of viable progeny was scored 48 h after tetrad dissection. n = 18 tetrads. (E) Representative images from live imaging of Mtw1-tdTomato and Ipl1-mNeonGreen at prophase I exit and anaphase I. All scale bars represent 2 μm. (F) Absence of Alk1 and Alk2 does not prevent Ipl1 localisation at kinetochores. Quantification of Ipl1-mNeonGreen signal intensity from (E) at prophase I exit. Boxplots represent interquartile ranges (IQR), and the middle line is the median. Whiskers extend to the furthest data points within 1.5 IQR from the 25th and 75th percentiles; data points outside this range are plotted individually as outliers; Mann-Whitney test. (G-I) The CPC is in close proximity to the outer kinetochore rather than Ctf19 at prophase I exit. Volcano plots presenting the mass spectrometry-identified biotinylated proteins pulled down by streptavidin-coated beads at prophase I exit in diploids expressing (G) Ctf19-miniTurboID *sli15ΔN*, compared to no tag; (H) Ctf19-miniTurboID *sli15ΔN* compared to Ctf19-miniTurboID; (I) sli15ΔN-miniTurboID compared to Sli15-miniTurboID. Diploids containing the Ndt80 block-release system were induced to undergo meiosis and released from arrest by β-estradiol addition at 6 h. Cells were harvested 30 min later. Data are from 3 biological replicates. Log2(fold change) between conditions is shown with corresponding p-values. Dashed line indicates log_2_(fold change) = |2|.

**Figure S4.**
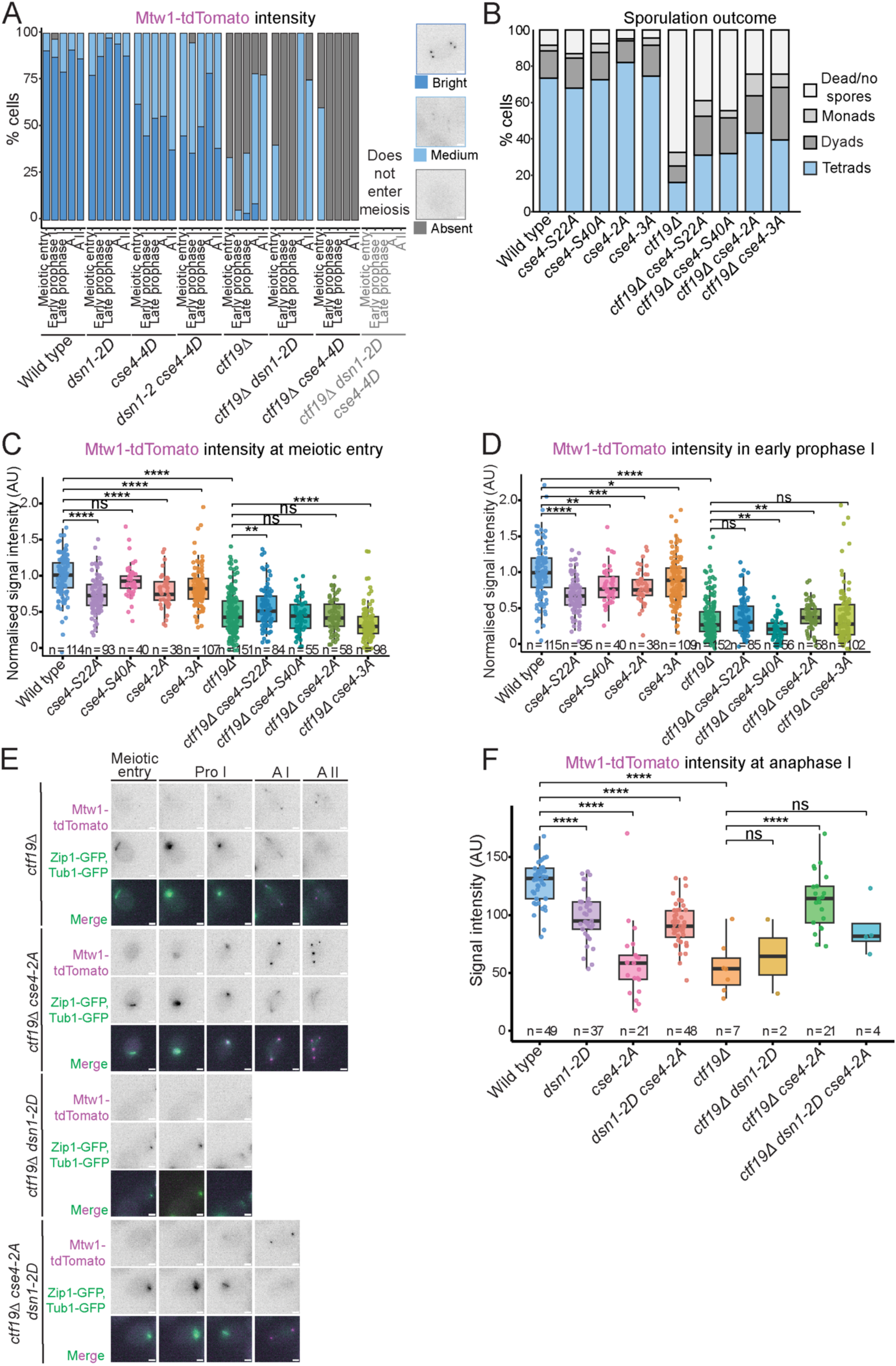
Dsn1 phosphorylation precludes kinetochore assembly in *ctf19Δ* cells despite the absence of Cse4 phosphorylation. Related to Figure 5. (A) Phosphorylation of Ipl1 target proteins in the kinetochore does not rescue *ctf19Δ* kinetochore assembly defect. Percentage of cells with ‘bright’, ‘medium’, or ‘absent’ Mtw1-tdTomato signal at each stage of meiosis, as determined by live imaging of Tub1-GFP and Zip1-GFP. Inset: representative images of each category. n = 100 cells per strain. (B) Preventing phosphorylation of Ipl1 target sites on Cse4 mildly improves sporulation efficiency of *ctf19Δ* cells. Percentage of cells forming tetrads, dyads, monads or not sporulating/dying. n = 200 cells scored for each strain. (C-D) Phosphomimetic mutations of *Cse4* do not improve kinetochore assembly in *ctf19Δ* cells. Quantification of Mtw1-tdTomato signal intensity from (Figure 5E) at (C) meiotic entry; (D) early prophase I. Meiotic stage determined by Zip1-GFP appearance. All scale bars represent 2 μm. In (C-D) and (F), boxplots represent interquartile ranges (IQR), and the middle line is the median. Whiskers extend to the furthest data points within 1.5 IQR from the 25th and 75th percentiles; data points outside this range are plotted individually as outliers; Mann-Whitney test. (E) Representative images from live imaging of Mtw1-tdTomato, Zip1-GFP, and Tub1-GFP. Meiotic stage determined by Zip1-GFP appearance and disappearance, and Tub1-GFP morphology. *ctf19Δ dsn1-S240D, S250D* cells did not progress past prophase I exit. *ctf19Δ cse4-2A dsn1-S240D, S250D* cells did not progress past anaphase I. (F) Dsn1 phosphorylation abrogates the rescue of *ctf19Δ* by *cse4-2A* at anaphase I. Quantification of Mtw1-tdTomato signal intensity from (E) at anaphase I.

**Figure S5.**
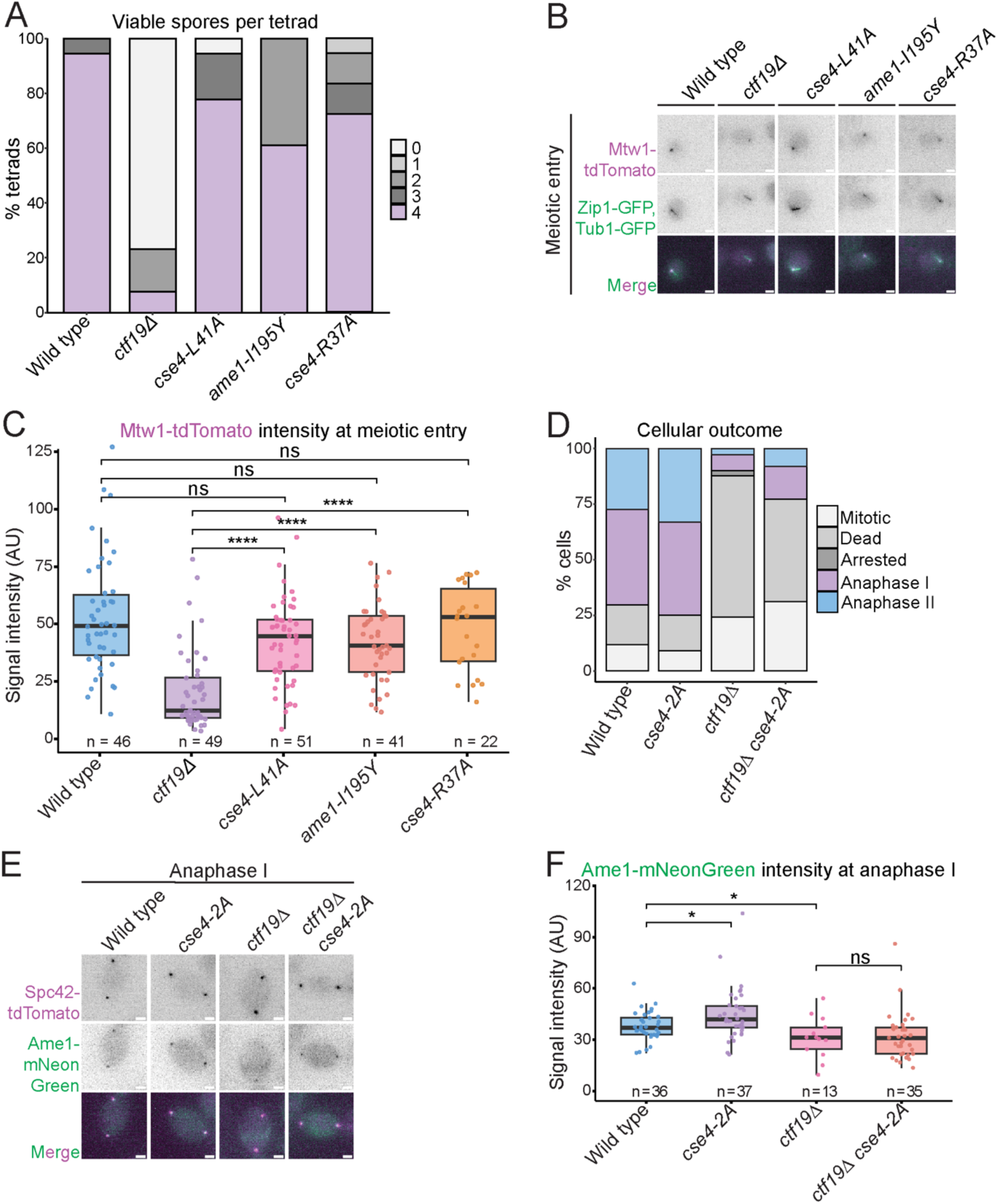
Okp1^CENP-Q^-Ame1^CENP-U^ are involved in two distinct kinetochore assembly pathways. Related to Figure 6. (A) Disrupting the Cse4^CENP-A^-Okp1^CENP-Q^-Ame1^CENP-U^ interaction does not impair spore viability as severely as *ctf19Δ.* The number of viable progeny was scored 48 h after tetrad dissection. n = 18 tetrads. (B) Representative images from live imaging of Mtw1-tdTomato, Zip1-GFP, and Tub1-GFP at meiotic entry. All scale bars represent 2 μm. C) Disrupting the Cse4^CENP-A^-Okp1^CENP-Q^-Ame1^CENP-U^ interaction does not prevent kinetochore assembly at meiotic entry. Quantification of Mtw1-tdTomato signal intensity from (B). In (C) and (F), boxplots represent interquartile ranges (IQR), and the middle line is the median. Whiskers extend to the furthest data points within 1.5 IQR from the 25th and 75th percentiles; data points outside this range are plotted individually as outliers; Mann-Whitney test. (D) Tagging Ame1 prevents meiotic progression in *ctf19Δ* and *ctf19Δ cse4-2A* cells. Percentage of cells reaching each outcome as indicated. (E) Representative images from live imaging of Spc42-tdTomato and Ame1-mNeonGreen at anaphase I. All scale bars represent 2 μm. (F) *cse4-2A* does not rescue Ame1-mNeonGreen signal in *ctf19Δ* cells. Quantification of Ame1-mNeonGreen signal intensity at anaphase I from (E).

## References

Akiyoshi B, Nelson CR & Biggins S (2013) The Aurora B Kinase Promotes Inner and Outer Kinetochore Interactions in Budding Yeast. Genetics 194: 785–789

Anedchenko EA, Samel-Pommerencke A, Tran Nguyen TM, Shahnejat-Bushehri S, Pöpsel J, Lauster D, Herrmann A, Rappsilber J, Cuomo A, Bonaldi T, et al (2019) The kinetochore module Okp1^CENP-Q^ /Ame1^CENP-U^ is a reader for N-terminal modifications on the centromeric histone Cse4^CENP-A^. EMBO J 38: e98991

Barton RE, Massari LF, Robertson D & Marston AL (2022) Eco1-dependent cohesin acetylation anchors chromatin loops and cohesion to define functional meiotic chromosome domains. eLife 11: e74447

Berchowitz LE, Gajadhar AS, Van Werven FJ, De Rosa AA, Samoylova ML, Brar GA, Xu Y, Xiao C, Futcher B, Weissman JS, et al (2013) A developmentally regulated translational control pathway establishes the meiotic chromosome segregation pattern. Genes Dev 27: 2147–2163

Bock LJ, Pagliuca C, Kobayashi N, Grove RA, Oku Y, Shrestha K, Alfieri C, Golfieri C, Oldani A, Dal Maschio M, et al (2012) Cnn1 inhibits the interactions between the KMN complexes of the yeast kinetochore. Nat Cell Biol 2012 146 14: 614–624

Bonner MK, Haase J, Swinderman J, Halas H, Miller Jenkins LM & Kelly AE (2019) Enrichment of Aurora B kinase at the inner kinetochore controls outer kinetochore assembly. J Cell Biol 218: 3237–3257

Borek WE, Vincenten N, Duro E, Makrantoni V, Spanos C, Sarangapani KK, de Lima Alves F, Kelly DA, Asbury CL, Rappsilber J, et al (2021) The Proteomic Landscape of Centromeric Chromatin Reveals an Essential Role for the Ctf19CCAN Complex in Meiotic Kinetochore Assembly. Curr Biol 31: 283–296.e7

Branon TC, Bosch JA, Sanchez AD, Udeshi ND, Svinkina T, Carr SA, Feldman JL, Perrimon N & Ting AY (2018) Efficient proximity labeling in living cells and organisms with TurboID. Nat Biotechnol 36: 880–887

Camahort R, Li B, Florens L, Swanson SK, Washburn MP & Gerton JL (2007) Scm3 Is Essential to Recruit the Histone H3 Variant Cse4 to Centromeres and to Maintain a Functional Kinetochore. Mol Cell 26: 853–865

Campbell CS & Desai A (2013) Tension sensing by Aurora B kinase is independent of survivin-based centromere localization. Nature 497: 118–121

Carlile TM & Amon A (2008) Meiosis I Is Established through Division-Specific Translational Control of a Cyclin. Cell 133: 280–291

Carroll CW, Silva MCC, Godek KM, Jansen LET & Straight AF (2009) Centromere assembly requires the direct recognition of CENP-A nucleosomes by CENP-N. Nat Cell Biol 11: 896–902

Celebic D, Polat I, Legros V, Chevreux G, Wassmann K & Touati SA (2024) Qualitative rather than quantitative phosphoregulation shapes the end of meiosis I in budding yeast. EMBO J 43: 1325–1350

Chen J, Liao A, Powers EN, Liao H, Kohlstaedt LA, Evans R, Holly RM, Kim JK, Jovanovic M & Ünal E (2020) Aurora B-dependent Ndc80 degradation regulates kinetochore composition in meiosis. Genes Dev 34: 209–225

Chen J, Tresenrider A, Chia M, McSwiggen DT, Spedale G, Jorgensen V, Liao H, van Werven FJ & Ünal E (2017) Kinetochore inactivation by expression of a repressive mRNA. eLife 6: e27417

Dai J, Sultan S, Taylor SS & Higgins JMG (2005) The kinase haspin is required for mitotic histone H3 Thr 3 phosphorylation and normal metaphase chromosome alignment. Genes Dev 19: 472–488

De Wulf P, McAinsh AD & Sorger PK (2003) Hierarchical assembly of the budding yeast kinetochore from multiple subcomplexes. Genes Dev 17: 2902–2921

Demichev V, Messner CB, Vernardis SI, Lilley KS & Ralser M (2020) DIA-NN: neural networks and interference correction enable deep proteome coverage in high throughput. Nat Methods 17: 41–44

Dendooven T, Zhang Z, Yang J, McLaughlin SH & Schwab J (2023) Cryo-EM structure of the complete inner kinetochore of the budding yeast point centromere. Sci Adv

Dendooven T, Zhang Z, Yang J, McLaughlin SH, Schwab J, Scheres SHW, Yatskevich S & Barford D Cryo-EM structure of the complete inner kinetochore of the budding yeast point centromere. Sci Adv 9: eadg7480

Deng S, Cai J, Harrison SC, Zhou H & Hinshaw SM (2023) Recognition of centromere-specific histone Cse4 by the inner kinetochore Okp1-Ame1 complex. EMBO Rep 24: e57702

Dimitrova YN, Jenni S, Valverde R, Khin Y & Harrison SC (2016) Structure of the MIND Complex Defines a Regulatory Focus for Yeast Kinetochore Assembly. Cell 167: 1014–1027.e12

Dudziak A, Pleuger R, Schmidt J, Hamm F, Tendulkar S, Jänen K, Vetter IR, Singh S, Fischböck J, Herzog F, et al (2025) The Spc105/Kre28 complex promotes mitotic error correction by outer kinetochore recruitment of Ipl1/Sli15. EMBO J 44: 3492–3520

Fernius J & Marston AL (2009) Establishment of Cohesion at the Pericentromere by the Ctf19 Kinetochore Subcomplex and the Replication Fork-Associated Factor, Csm3. PLoS Genet 5: e1000629

Fischböck-Halwachs J, Singh S, Potocnjak M, Hagemann G, Solis-Mezarino V, Woike S, Ghodgaonkar-Steger M, Weissmann F, Gallego LD, Rojas J, et al (2019) The COMA complex interacts with Cse4 and positions Sli15/Ipl1 at the budding yeast inner kinetochore. eLife 8: e42879

Furuyama S & Biggins S (2007) Centromere identity is specified by a single centromeric nucleosome in budding yeast. Proc Natl Acad Sci 104: 14706–14711

García-Rodríguez LJ, Kasciukovic T, Denninger V & Tanaka TU (2019) Aurora B-INCENP Localization at Centromeres/Inner Kinetochores Is Required for Chromosome Bi-orientation in Budding Yeast. Curr Biol 29: 1536–1544.e4

Gascoigne KE, Takeuchi K, Suzuki A, Hori T, Fukagawa T & Cheeseman IM (2011) Induced Ectopic Kinetochore Assembly Bypasses the Requirement for CENP-A Nucleosomes. Cell 145: 410–422

Ghosh SK, Poddar A, Hajra S, Sanyal K & Sinha P (2001) The IML3/MCM19 gene of Saccharomyces cerevisiae is required for a kinetochore-related process during chromosome segregation. Mol Genet Genomics 265: 249–257

Greco M, Klim J & De Wulf P (2026) Keeping it centered: decoding the activities that regulate yeast histone H3 variant Cse4 and confine it to centromeres. Front Cell Dev Biol 14: 1782273

Hamilton GE, Helgeson LA, Noland CL, Asbury CL, Dimitrova YN & Davis TN (2020) Reconstitution reveals two paths of force transmission through the kinetochore. eLife 9: e56582

Hayashi A, Asakawa H, Haraguchi T & Hiraoka Y (2006) Reconstruction of the kinetochore during meiosis in fission yeast Schizosaccharomyces pombe. Mol Biol Cell 17: 5173–5184

Hinshaw SM & Harrison SC (2018) Kinetochore Function from the Bottom Up. Trends Cell Biol 28: 22–33

Hinshaw SM & Harrison SC (2019) The structure of the Ctf19c/CCAN from budding yeast. eLife 8: e44239

Hinshaw SM, Makrantoni V, Harrison SC & Marston AL (2017) The Kinetochore Receptor for the Cohesin Loading Complex. Cell 171: 72–84.e13

van Hooff JJ, Tromer E, van Wijk LM, Snel B & Kops GJ (2017) Evolutionary dynamics of the kinetochore network in eukaryotes as revealed by comparative genomics. EMBO Rep 18: 1559–1571

Hori T, Shang W-H, Takeuchi K & Fukagawa T (2013) The CCAN recruits CENP-A to the centromere and forms the structural core for kinetochore assembly. J Cell Biol 200: 45–60

Hornung P, Troc P, Malvezzi F, Maier M, Demianova Z, Zimniak T, Litos G, Lampert F, Schleiffer A, Brunner M, et al (2014) A cooperative mechanism drives budding yeast kinetochore assembly downstream of CENP-A. J Cell Biol 206: 509–524

Killinger K, Böhm M, Steinbach P, Hagemann G, Blüggel M, Jänen K, Hohoff S, Bayer P, Herzog F & Westermann S (2020) Auto-inhibition of Mif2/CENP-C ensures centromere-dependent kinetochore assembly in budding yeast. EMBO J 39

Knop M, Siegers K, Pereira G, Zachariae W, Winsor B, Nasmyth K & Schiebel E (1999) Epitope tagging of yeast genes using a PCR-based strategy: more tags and improved practical routines. Yeast 15: 963–972

Koch LB & Marston AL (2025) The functional organisation of the centromere and kinetochore during meiosis. Curr Opin Cell Biol 94: 102486

Koch LB, Spanos C, Kelly V, Ly T & Marston AL (2024) Rewiring of the phosphoproteome executes two meiotic divisions in budding yeast. EMBO J 43: 1351–1383

Longtine MS, Mckenzie Iii A, Demarini DJ, Shah NG, Wach A, Brachat A, Philippsen P & Pringle JR (1998) Additional modules for versatile and economical PCR-based gene deletion and modification in Saccharomyces cerevisiae. Yeast 14: 953–961

Makrantoni V, Ciesiolka A, Lawless C, Fernius J, Marston A, Lydall D & Stark MJR (2017) A functional link between bir1 and the Saccharomyces cerevisiae Ctf19 kinetochore complex revealed through quantitative fitness analysis. G3 Genes Genomes Genet 7

Malvezzi F, Litos G, Schleiffer A, Heuck A, Mechtler K, Clausen T & Westermann S (2013) A structural basis for kinetochore recruitment of the Ndc80 complex via two distinct centromere receptors. EMBO J 32: 409–423

Meyer RE, Chuong HH, Hild M, Hansen CL, Kinter M & Dawson DS (2015) Ipl1/Aurora-B is necessary for kinetochore restructuring in meiosis I in Saccharomyces cerevisiae. Mol Biol Cell 26: 2986–3000

Meyer RE, Kim S, Obeso D, Straight PD, Winey M & Dawson DS (2013) Mps1 and Ipl1/Aurora B Act Sequentially to Correctly Orient Chromosomes on the Meiotic Spindle of Budding Yeast. Science 339: 1071–1074

Miller MP, Ünal E, Brar GA & Amon A (2012) Meiosis I chromosome segregation is established through regulation of microtubule-kinetochore interactions. eLife 2012

Monje-Casas F, Prabhu VR, Lee BH, Boselli M & Amon A (2007) Kinetochore Orientation during Meiosis Is Controlled by Aurora B and the Monopolin Complex. Cell 128: 477–490

Ng TM, Waples WG, Lavoie BD & Biggins S (2009) Pericentromeric sister chromatid cohesion promotes kinetochore biorientation. Mol Biol Cell 20: 3818–3827

Ortiz J, Stemmann O, Rank S & Lechner J (1999) A putative protein complex consisting of Ctf19, Mcm21, and Okp1 represents a missing link in the budding yeast kinetochore. Genes Dev 13: 1140–1155

Popchock AR, Hedouin S, Mao Y, Asbury CL, Stergachis AB & Biggins S (2025) Stable centromere association of the yeast histone variant Cse4 requires its essential N-terminal domain. EMBO J 44: 1488–1511

Przewloka MR, Venkei Z, Bolanos-Garcia VM, Debski J, Dadlez M & Glover DM (2011) CENP-C is a structural platform for kinetochore assembly. Curr Biol 21: 399–405

Rabitsch KP, Petronczki M, Javerzat J-P, Genier S, Chwalla B, Schleiffer A, Tanaka TU & Nasmyth K (2003) Kinetochore recruitment of two nucleolar proteins is required for homolog segregation in meiosis I. Dev Cell 4: 535–548

Rappsilber J, Mann M & Ishihama Y (2007) Protocol for micro-purification, enrichment, pre-fractionation and storage of peptides for proteomics using StageTips. Nat Protoc 2: 1896–1906

Samel A, Cuomo A, Bonaldi T & Ehrenhofer-Murray AE (2012) Methylation of CenH3 arginine 37 regulates kinetochore integrity and chromosome segregation. Proc Natl Acad Sci 109: 9029–9034

Schleiffer A, Maier M, Litos G, Lampert F, Hornung P, Mechtler K & Westermann S (2012) CENP-T proteins are conserved centromere receptors of the Ndc80 complex. Nat Cell Biol 14: 604–613

Sears RM, May DG & Roux KJ (2019) BioID as a Tool for Protein-Proximity Labeling in Living Cells. In Enzyme-Mediated Ligation Methods, Nuijens T & Schmidt M (eds) pp 299–313. New York, NY: Springer New York

Shevchenko A, Wilm M, Vorm O & Mann M (1996) Mass Spectrometric Sequencing of Proteins from Silver-Stained Polyacrylamide Gels. Anal Chem 68: 850–858

Shukla S, Bhattacharya A, Sehrawat P, Agarwal P, Shobhawat R, Malik N, Duraisamy K, Rangan NS, Hosur RV & Kumar A (2024) Disorder in CENP-ACse4 tail-chaperone interaction facilitates binding with Ame1/Okp1 at the kinetochore. Structure 32: 690–705.e6

Turner NN, Zhang Z, Yang J, Muir KW, McLaughlin SH, Morgan T & Barford D (2026) Assembly and phosphoregulatory mechanisms of the budding yeast outer kinetochore KMN complex. J Cell Biol 225: e202506015

Vincenten N, Kuhl L-M, Lam I, Oke A, Kerr AR, Hochwagen A, Fung J, Keeney S, Vader G & Marston AL (2015) The kinetochore prevents centromere-proximal crossover recombination during meiosis. eLife 4: e10850

Vinod A, Turner M, Webb S & Marston AL (2026) Multiple pathways couple kinetochore orientation to the meiotic spindle cycle. doi:10.64898/2026.04.17.719180 [PREPRINT]

Wang F, Dai J, Daum JR, Niedzialkowska E, Banerjee B, Stukenberg PT, Gorbsky GJ & Higgins JMG (2010) Histone H3 Thr-3 Phosphorylation by Haspin Positions Aurora B at Centromeres in Mitosis. Science 330: 231–235

Xiao H, Wang F, Wisniewski J, Shaytan AK, Ghirlando R, FitzGerald PC, Huang Y, Wei D, Li S, Landsman D, et al (2017) Molecular basis of CENP-C association with the CENP-A nucleosome at yeast centromeres. Genes Dev 31: 1958–1972

Yan K, Yang J, Zhang Z, McLaughlin SH, Chang L, Fasci D, Ehrenhofer-Murray AE, Heck AJR & Barford D (2019) Structure of the inner kinetochore CCAN complex assembled onto a centromeric nucleosome. Nature 574: 278–282

Yoon H-J & Carbon J (1999) Participation of Bir1p, a member of the inhibitor of apoptosis family, in yeast chromosome segregation events. Proc Natl Acad Sci 96: 13208–13213

